# Multi-year study on the effects of elevated CO_2_ in mature oaks unravels subtle metabolic adjustments but stable biotic stress resistance

**DOI:** 10.1101/2025.05.03.652050

**Authors:** Rosa Sanchez-Lucas, Mark Raw, Alisha Datta, Katie Hawkins, Deanne Brettle, Emma Ann Platt, Sami Ullah, Kris Hart, Carolina Mayoral, Moritz Stegner, Ilse Kranner, Scott A. L. Hayward, Victoria Pastor, A. Rob MacKenzie, Estrella Luna

## Abstract

- Rising atmospheric CO_2_ levels are predicted to influence forest health directly and indirectly, yet the long-term effects of elevated CO_2_ (eCO_2_) on mature trees in natural ecosystems remain poorly understood. Understanding how eCO_2_ affects susceptibility to biotic stress and alters leaf metabolism is critical for predicting forest responses to climate change.
- We examined the effects of eCO_2_ (+150 ppm) on 180-year-old *Quercus robur* at the Birmingham Institute of Forest Research (BIFoR) Free Air CO_2_ Enrichment (FACE) facility. From 2016 (pre-treatment) to 2024 (year 8 of enrichment), we monitored natural powdery mildew infection and insect herbivory, alongside targeted and untargeted metabolomic profiling of leaf material collected across the growing season.
- While seasonal patterns and an overall decline in PM and herbivory were observed, no consistent differences in biotic stress incidence emerged due to eCO_2_. Metabolomic data revealed subtle but widespread shifts, especially in amino acid, CoenzymeA, and redox pathways.
- These results suggest that although eCO_2_ drives extensive metabolic changes, it does not alter biotic stress resistance in mature oaks. Instead, eCO_2_ appears to promote physiological plasticity that may shape future responses to combined environmental stressors. These insights offer a valuable reference point for interpreting long-term ecosystem dynamics.

## Introduction

Atmospheric carbon dioxide (CO_2_) levels are at their highest point in over 3 million years (Keeling, 2017). In the last century, they have increased from 285 parts per million (ppm) just before the industrial revolution, to over 420 ppm at present (Lan, 2025). By 2100, predictions estimate that CO_2_ levels will rise to between 540 and 970 ppm (Song *et al*., 2022). Elevated CO_2_ (eCO_2_) levels impact on global climate, resulting in a rise in temperature and more frequent and more extreme weather events such as floods or droughts (Baker *et al*., 2018; IPCC, 2023). For instance, the 10 warmest years on record in the United Kingdom have all occurred since 2002 (Met Office, 2023). To prevent further damage, countries have recognised the importance of reducing CO_2_ emissions, however, removing and storing current CO_2_ from the atmosphere has also become of extreme importance. Globally, the land carbon sink, dominated by forest systems, capture up to 30% of the total anthropogenic CO_2_ emitted into air per year (Friedlingstein *et al*., 2025). Nevertheless, the importance of forests extends way beyond their CO_2_ capture potential. For instance, forests can be highly varied and productive ecosystems becoming hubs of biodiversity. The world is witnessing a rapid reduction in species diversity and estimates of around 1 million species are at threat of extinction (Díaz, 2019). Moreover, forests are also important for the global economy by supporting an estimated 13.2 million jobs and producing a gross value of $600 billion per year (Food and Agriculture Organization of the United Nations, 2014). Therefore, tree planting and forest protection can mitigate the impact of global challenges and specifically contribute to activities to control and reduce levels of atmospheric CO_2_ (IPCC, 2023; Norby *et al*., 2024).

Oak trees (Quercus spp.) are a vital component of forests around the world and, in the UK, *Q. robur* and *Q. petrea* and their hybrids are the 3rd most common broad-leaved tree with over 121 million specimens (Quine *et al*., 2019) and oak is now an integral part of tree planting strategies (Whittet *et al*., 2019). Given the importance of oak forests, it is crucial to consider any threats they may face. These include biotic stresses such as fungal pathogens and insect pests. One of those fungal diseases is powdery mildew (PM). Different biotrophic fungal pathogens cause PM disease, the most common ones being *Erysiphe alphitoides* and *E. quercicola,* which in oaks is characterised mostly by a white powder covering the leaf surface causing tissue decay. PM disease occurs annually at varying levels in oak forests (Marçais *et al*., 2017). The disease reduces photosynthetic efficiency and carbon assimilation whilst increasing the rates of transpiration (Marçais & Desprez-Loustau, 2014). This results in a remarkably high seedling mortality rate (Marçais *et al*., 2009; Sanchez-Lucas *et al*., 2023), which turns PM into a bottleneck to woodland regeneration and expansion. Importantly, whereas mature oak trees have been observed to mitigate well the impacts of the disease under ‘normal’ environmental circumstances (i.e. without stress) (Mayoral *et al*., 2023), reports indicate that in association with other stresses, such as drought and high temperatures, it can result in the decline and death of mature oak trees (Lonsdale, 2016). Understanding this increased vulnerability to PM is of great importance given the increasing frequency of heat wave and drought events under predicted climate change (Marçais & Desprez-Loustau, 2014). This is because the disease could not only threaten woodland regeneration, but also the conservation of established mature, and highly biodiverse, woodland ecosystems.

Insect infestations also pose a significant threat to oak trees. Herbivores, ranging from insects to mammals, feed on various parts of oak trees, including leaves, buds, and acorns. The loss of foliage due to herbivory can reduce the tree’s capacity to photosynthesise, limiting its energy production consequently stunting growth (Alalouni *et al*., 2014). Additionally, repeated defoliation over several years can lead to a decline in overall tree health, affecting reproduction and making the tree more vulnerable to mortality (Kulman, 1971). While some oak species can partially recover from defoliation events, especially if they occur early in the growing season, prolonged or severe infestations can have long lasting ecological consequences, impacting not only the affected trees but also the entire forest ecosystem (Rieske & Dillaway, 2008). A particularly damaging insect that has dire effects on the oak tree is the European Winter Moth, (*Operophtera brumata*). These insects feed on oak leaves during their larval stage, often defoliating entire trees in the process (Roland, 1995). Severe infestations can weaken oak trees, making them more susceptible to other stressors such as disease. Considering these complex challenges, comprehensive forest protection and conservation efforts are imperative. Understanding interactions between pathogen and herbivore stress is also important. Damage caused by an initial infestation (pest or pathogen) can subsequently lead to a stressed tree becoming more vulnerable to another stressor (Teshome *et al*., 2020). For example, extensive early season insect defoliation on *Q. robur* can promote second flush of new growth (‘lammas shoots’) that are then infected by oak powdery mildew (Glazebrook, 2005; Marçais & Desprez-Loustau, 2014). These sequential and potentially compounding biotic stressors on the host plant, can in turn trigger different plant defence signalling pathways, leading to crosstalk between pathways and antagonistic or facilitative interactions between attackers (Erb, 2012). Elevated CO_2_ may complicate this interaction yet further with evidence of both increased herbivory under eCO_2_, due to the potentially reduced nutritional value of leaves with higher C:N ratios (Robinson, 2012), but also examples of reduced herbivory under eCO_2_ allowing more resource for plants to increase the production of defence compounds (Ryan *et al*., 2010).

Whilst CO_2_ is often villainised for its impact in climate change, it is also a vital commodity for all photosynthetic life. The exposure to eCO_2_ particularly in young plants has been shown to promote growth driven by increased photosynthesis and increased nitrogen use efficiency (Norby *et al*., 1986). Working in the natural conditions of the Birmingham Institute of Forest Research (BIFoR) Free Air CO_2_ Enrichment (FACE) facility, Gardner *et al*. (2022) found sustained photosynthetic enhancement over 3 years of eCO_2_ exposure in mature oak trees. Whereas this could be seen as a positive CO_2_ fertilization effect, there is evidence that eCO_2_ can impact other processes such as plant defence mechanisms (Smith & Luna, 2023). However, the effects of eCO_2_ in plant defence are complex and can result in highly contrasting resistance phenotypes. For instance, in soybean (*Glycine max*), eCO_2_ was shown to have a positive effect in the resistance against stem canker and this phenotype was linked to higher levels of the key defence hormone salicylic acid (SA) and phytoalexins (Braga *et al*., 2006). In contrast, other studies have detected increased susceptibility to different pathogens (Lake & Wade, 2009; Lessin & Ghini, 2009; Pugliese *et al*., 2010), including PMs in *Arabidopsis thaliana*, soybean and grape. These disease phenotypes were linked to numerous factors including increased stomatal density, as well as alterations in metabolites such as SA and Jasmonic acid (JA) accumulation (Kazan, 2018). Intriguingly, eCO_2_ substantially dampened jasmonate signalling in Lucerne (*Mendicago sativa*) post Cotton Bollworm (*Helicoverpa armigera*) attack, resulting in significantly lower JA concentrations in attacked plants grown under eCO_2_ (Johnson *et al*., 2020). This reduction in defence signalling coincided with a noteworthy increase in the Cotton Bollworm’s relative growth rates when consuming eCO_2_-grown Lucerne. We have previously reported that oak seedlings grown under eCO_2_ conditions showed greater susceptibility to PM infection than their counterparts grown under ambient CO_2_ (aCO_2_, Sanchez-Lucas et al 2023). This phenotype has been linked to potential trade-offs between growth and defence under eCO_2_. Importantly, exposing plants to elevated levels of CO_2_ leads to a nuanced impact on their nutritional profile. The research highlights positive outcomes, including an increase in concentrations of beneficial components such as fructose, glucose, total antioxidant capacity but also with a significant decline in total nitrogen concentrations (Dong *et al*., 2018). The nutritional quality of the host plant can significantly impact rates of herbivory (Hamilton *et al*., 2004), and thus herbivore growth or reproduction rates (Crowley, 2021), although these outcomes are often species specific. Therefore, it is clear that eCO_2_ can negatively impact diseases and pests in different plant species. Nevertheless, these experiments were done in highly controlled growth conditions and with levels of CO_2_ that will unlikely be achieved on our planet. However, a study using realistic concentrations of CO_2_ and in real environmental conditions in the first three years of eCO_2_ at Birmingham Institute of Forest Research (BIFoR) Free Air CO_2_ Enrichment (FACE) revealed no overall change in herbivory damage from leaf litter samples (Roberts *et al*., 2022). Moreover, the molecular mechanisms responsible for the impact of eCO_2_ in oak immunity and metabolism remain unknown. Further mechanistic studies are therefore needed to unravel the effects of eCO_2_, and these need to be undertaken in natural forest systems and not just in a lab.

Here, we aim to assess the global impact of eCO_2_ (ambient +150 ppm) in a mature temperate forest on the susceptibility of mature oak (*Q. robur*) trees to PM and insect damage. We hypothesise that eCO_2_ will result in altered resistance phenotypes, most likely enhancing susceptibility. Moreover, we also hypothesise that eCO_2_ will result in specific biochemical profiles in the exposed trees. Specifically, trees may adjust the composition of their photosynthetic metabolites under eCO_2_, to optimize light capture and energy conversion, which could lower levels of effective stress avoidance mechanisms, preventing the production of defensive responses such as reactive oxygen species (ROS). To test these hypotheses, we monitored natural PM infections and insect damage for nine years in mature oak trees. Moreover, we performed a targeted analysis of photosynthesis-related metabolites and an untargeted metabolome analysis at different points of the growth season to identify potential mechanisms associated with different susceptibility outcomes due to eCO_2_ exposure. Our results provide the most detailed insight to date of how mature, long-lived, plants respond metabolically to multi-year eCO_2_ and how this may impact biotic stress. These data could provide a foundation from which to develop guidelines for forest management practices and policy implementation striving to enhance oak woodland conservation.

## Methods

### 1. Experimental Free-Air CO_2_ Enrichment (FACE) facility

All experiments described in this work were done at the Free Air Carbon Enrichment (FACE) facility of the Birmingham Institute of Forest Research (BIFoR). BIFoR-FACE is a unique facility for forest climate change research where six patches of an unmanaged 180 year-old oak forest are exposed to ambient (aCO_2_; n=3) and elevated (eCO_2_; n=3) levels of CO_2_. The facility consists of six approximately circular ‘arrays’ (As) of towers that extend to top-of-canopy and encompass a patch of woodland approximately 30 m in diameter. Three of these arrays (A2, A3 and A5) blow ambient air into the prevailing winds and the other three arrays (A1, A4 and A6) mix CO_2_ into the air flow. The eCO_2_ target treatment is to elevate the atmosphere in the treatment arrays by an additional 150 ppm of CO_2_ over the daylight period of each day in the deciduous growing season. The experiment was switched on in April 2017 and will continue to run during each growing season (April-October inclusive) until at least 2031 (Hart *et al*., 2020; MacKenzie *et al*., 2021; Gardner *et al*., 2022). The ambient and enriched CO_2_ levels for each growing season can be found in Supplementary Table 1.

### 2. Experimental procedures for the study of powdery mildew infection and insect infestation

#### 2.1. Selected oak trees

All trees within the BIFoR-FACE facility with a diameter >10 cm, at 1.3 m stem height, have been assigned a four-digit identifier. Leaf materials for analysis were gathered from two selected mature trees per array. Selected trees are located in the middle of each array and have been monitored since 2016 to date. Selected trees correspond with the following labelling: Array 1 (A1) 8446 and 8437; Array 2 (A2) 8749 and 8673; Array 3 (A3) 8343 and 9301; Array 4 (A4) 6632 and 6622; Array 5 (A5) 6387 and 6382; and Array 6 (A6) 3752 and 5846. Due to experimental requirements from the larger institute’s research community, during the year of 2020, A4 swapped the tree 6622 for tree 6621 and from 2021 onwards, the three trees 6632, 6622 and 6621 were included in different analyses.

#### 2.2. Leaf sampling

Selected mature trees from the different arrays were used for this analysis. Array 4 material was obtained from trees 6632 and 6622 from 2016 to 2019, from 6632 and 6621 in 2020, and from 6632, 6622 and 6621 in 2021, 2022, 2023 and 2024. Oak leaves were collected monthly during the photosynthetically active period. PM disease symptoms could be observed on the leaves from June from 2016 to 2021 and May in 2022, 2023 and 2024. PM was scored (Section 2.4) until September in all years. Insect damage could be observed from May and it was scored (Section 2.5) until September in all years. Arborists ascended the selected trees and collected three terminal branch sections of ∼20 cm with an average of 20 leaves per branch. One branch was brought down from each of the three canopy levels: top, middle and bottom. Canopy levels were determined by climbers using a laser distance finder (Leica DISTRO D2) to estimate the heights of each sampled branch, which have been consistent throughout the years. Subsequently, leaves were transferred to sample bags and stored at -20°C until processing.

#### 2.3. Leaf processing for image analysis

Stored leaves from 2016, 2017, 2018, 2019, 2021 were processed during the summer of 2021. Due to the larger institute’s research community requirements, the year 2020 was processed in the Autumn of 2020. Leaves from 2022, 2023 and 2024 were processed during the Autumn of their collection. Leaves were defrosted at room temperature using paper towel to remove excess water from the leaf surfaces. Between six to 15 leaves per collection point were placed adaxial side down on a light blue paper sheet (for background) containing a Helix L16 ruler, scanned at 300dpi using a CanoScan LiDE 220 scanner and saved in .tif format. Preparation and scanning of the 2020 leaves were performed in a similar manner except that a white graph (gridded) background was used.

#### 2.4. Quantification of powdery mildew disease

Scanned images were employed to quantify powdery mildew disease. Using FIJI software (Schindelin *et al*., 2012) we developed a script consisting of a 4-step pipeline: 1) removal of the background, 2) quantification of the leaf area, 3) masking out of senesced tissue (ie. brown and necrotic lesions); and 4) PM assessment through automatic quantification of white mycelium symptoms on the leaves. The analysis pipeline with images and macro was developed to standardise, process, and quantify powdery mildew coverage on oak leaves by cropping the image to remove irrelevant areas, detecting and thresholding powdery mildew areas using HSB (i.e., hue, saturation, brightness) colour space, creating a binary mask for clear segmentation and measuring powdery mildew coverage and storing results. Steps 1 and 2 also managed the removal and quantification of missing tissue due to insect damage. The analysis pipeline with image examples and scripts are available at https://github.com/PlantPriming/MatureOakCO2.

#### 2.5. Quantification for insect damage

All scanned leaves underwent a detailed scoring process, focusing on assessing the extent of chewing insect damage (i.e., herbivores that make holes on the leaves). The scoring system comprising six levels (i.e., categories: *d* = 0,1…*d_max_*; *d_max_*=5) was applied consistently across all sampled leaves. The categories were assigned depending on the amount of holes present on the leaves and the percentage of leaf area with insect damage (Supplementary Figure 1). The categories were: 0- indicating no damage, corresponding with 0% leaf area with damage; 1- small holes, corresponding with <10% leaf area with damage; 2- larger holes, corresponding with 10- 25% leaf area with damage; 3- significant holes, corresponding with 26-50% leaf area with damage; 4- significant leaf damage, corresponding with 51-75% leaf area with damage; and 5- near complete/ complete leaf damage, corresponding with 75%< leaf area with damage. Here, "holes" refer to areas where leaf tissue is missing, typically due to external chewing herbivory, and can occur both centrally and along the leaf margins. Image examples for category classification can be found in Supplementary Figure 1. Areas of brown tissue and other obvious insect damage, such as chlorosis or leaf mines, were excluded from the assessment. A damage index, *D*, was calculated for each image of *N* leaves, as the weighted average of damage scores in the image, expressed as a percentage of the maximum damage score, *d_max_*:

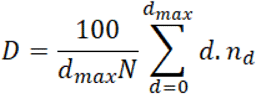

where *nd* is the number of leaves with damage score, *d*, and *d_max_* = 5.

#### 2.6 Statistical analysis and plots

Data underwent analysis using the coding program Rstudio. All scripts and commands are included in https://github.com/PlantPriming/MatureOakCO2. Statistical analyses were conducted utilising RStudio (version 1.4.1103) packages including Rmisc, dplyr, ggplot2, ggpubr, and effectsize. Statistical analyses were performed to assess the effect of eCO_2_ levels. Group-wise comparisons were conducted using non-parametric tests. Kruskal-Wallis tests were used to identify statistically significant differences among all groups. Wilcoxon rank-sum tests were used to assess pairwise comparisons. Cohen’s d was computed to quantify the effect size of the observed differences between different CO_2_ conditions. Cohen’s d values were interpreted following conventions for effect size interpretation. The data was visualised into box plots and line plots to illustrate the distribution scores across CO_2_ conditions, canopy layers, years and months. Benjamini-Hochberg p value adjustment was used on data represented as box plots as false discovery rate (FDR) correction.

### 3. Targeted identification of photosynthetic metabolites

#### 3.1 Leaf sample collection

Leaf samples were collected in August 2021 as described in section 2.1. From the leaves brought down from the mid-canopy (∼20) of the selected trees by arborists (i.e. six trees for aCO_2_, seven trees for eCO_2_ - two trees per array apart from A4 where 3 trees were sampled), six to 10 leaves were placed in a 50 ml tube and snapped frozen in the field by putting the tubes into a pre-cooled dry shipper (Statebourne Biotrek 10) following the dry shipper’s manufacturer specifications. Dry shipper was then transported to the lab and tubes were stored in the -80°C freezer until processing.

#### 3.2 Identification of low-molecular-weight thiols and disulfides

Low-molecular-weight (LMW) thiols and disulfides were extracted from 20 mg of finely ground leaf material in 1 mL of ice-cold 0.1 M HCl using a TissueLyser II (Qiagen) with two 5 mm glass beads at a frequency of 30 Hz for 2 minutes, and were analysed according to Bailly and Kranner (2011). 80 mg of polyvinylpolypyrrolidone (PVPP) was added to the leaf powder prior to extraction to bind phenolic compounds reacting with LMW thiols during extraction. Following extraction, the homogenate was centrifuged at 29,000 g for 20 minutes at 4°C. The supernatant was divided into two aliquots: a 120 μL aliquot for the quantification of LMW thiols + disulphides and a 400 μL aliquot for quantification of disulfides. In the first aliquot, 30 μL of 3 mM dithiothreitol (DTT) was added to reduce disulfides to thiols. In the second aliquot, thiols were blocked with N-ethylmaleimide (NEM), followed by removal of NEM and NEM-adducts by extracting five times with toluene prior to reduction with DTT, ensuring that only disulfides were quantified. In both cases, the thiols were subsequently derivatised with 20 μL of 15 mM monobromobimane, enabling fluorescence-based detection (excitation: 380 nm, emission: 480 nm). Separation and quantification of cysteine, γ-glutamyl-cysteine (γ-Glu-Cys), cysteinyl-glycine (Cys-Gly), and glutathione (GSH) were conducted using a reversed-phase Agilent 1260 Infinity II system with an Agilent 1290 Infinity II FLD-detector and a Phenomenex, Kinetex 5µm C18 100A, LC Column. Solvent A consisted of water:acetic acid, 99.75:0.25, v/v, pH 3.85. and solvent B was 100% methanol. Using a flow rate of 1 mL min^-1^ the run was started with 12% solvent B, followed by a gradient to 23% B over 11 minutes. Between 11.1 and 16 min the column was washed with 100% B, followed by equilibration with 12% B for 4 min before the next injection. The concentrations of LMW thiols and their corresponding disulphides were calculated using external standards, and thiols were calculated by subtracting disulphides concentrations from the concentrations of thiols + disulphides, as previously described by Bailly and Kranner (2011).

#### 3.3. Identification of photosynthetic pigments and tocochromanols

Pigments and tocochromanols were separated and quantified as described in Schausberger *et al*. (2019) and Buchner *et al*. (2017). Compounds were extracted from 20 mg of ground leaf material by adding 800 µL of methanol to each sample and homogenized with a TissueLyser II (Qiagen), with two 5 mm glass beads, for 5 minutes at 30 Hz, using racks that had been pre-cooled at −80 °C for 20 min. After homogenization, the samples were centrifuged at 29,000 g for 40 minutes at 4°C. A total of 400 µL of the supernatant was transferred into HPLC vials and 10 µL of the extract was injected into an Agilent 1100 HPLC system (Agilent Technologies) equipped with a LiChroCART® column (LiChrospher 100 RP-18, 125 × 4 mm, 5.0 µm particle size; Merck).

Tocochromanols were analysed using a fluorescence detector with an excitation wavelength of 295 nm and an emission wavelength of 325 nm. Simultaneously, the full-spectrum absorbance of pigments was measured using a diode-array detector at a wavelength of 440 nm. The mobile phase consisted of solvent A (acetonitrile:methanol, 74:6) and solvent B (methanol:hexane, 5:1), applied in a gradient over a 20-minute run at a flow rate of 1.0 mL min⁻¹. The run started at 0% B for the first 4 minutes, followed by a gradient to 100% B over the next 5 minutes, which was maintained for another 9 minutes.

#### 3.4 Statistics and plots

Plots and statistical analysis was conducted utilising RStudio (version 1.4.1103) packages ggplot2, Rmisc, dplyr, ggpur, and effectsize. Statistical significance was assessed using a Tukey test at a significance level of p value below 0.05. All scripts and commands are included in https://github.com/PlantPriming/MatureOakCO2.

### 4. Untargeted metabolomic analysis

#### 4.1 Leaf sample collection

Leaf samples were collected in May to September 2021 exactly as described in sections 2.1 and 3.1.

#### 4.2. Sample processing and metabolite extraction

Leaf material from mature trees and seedlings were homogenised using a pestle and mortar with liquid nitrogen and lyophilised. Metabolites were extracted using the protocol as described in (Manresa-Grao *et al*., 2022). Briefly, 1 mL of 30% methanol (MeOH) HPLC grade supplemented with 0.01% of HCOOH was added to 30 mg powdered lyophilised tissue. Tissue was incubated for 30 min on ice and then centrifuged for 30 min at 19000 g at 5°C. The supernatant was filtered using a 0.2 µm cellulose filter (Phenomenex). Metabolite extracts were stored at -80C until analysis.

#### 4.3. Mass spectrometry by LC-ESI full scan mass spectrometry in tandem (Q-TOF instrument)

A 20 µL aliquot of 1:3 diluted sample was injected into the UPLC in positive electrospray ionisation (ESI+) and negative (ESI-) ion modes for electrospray ionisation. A reversed-phase Acquity UPLC system (Waters, Milford, MA, United States) using a C-18 column (Kinetex 2.6 μm EVO C18 100 ÅLC Column 50 x 2.1 mm) with a gradient of MeOH and H_2_O supplemented with 0.01% HCOOH coupled to a hybrid quadrupole time-of-flight instrument (QTOF MS Premier) was used to elute and detect metabolites.

#### 4.4. Bioinformatic processing of metabolomic signals and metabolites identification

The Masslynx 4.1 (Masslynx 4.1, Waters) Databridge tool was used to transform data from the. raw format to .cdf files. Detected signals were processed using XCMS R scripts (Smith *et al*., 2006) to assign peaks, grouping, and signal corrections. Metabolite relative intensities were obtained based on the normalised peak area units relative to the dry weight of leaf sample. Positive and negative mode datasets were adduct adjusted and combined using MarVis (Version 6.7.3) filter software. The combined file was exported in .csv format and statistical tests were performed using the MetaboAnalyst (Version 6.0) platform. Data was normalised by sum and data scaling by Pareto scaling per month. Principal Component Analysis (PCAs) were performed for each month. Focussing on the effect of eCO_2_, we performed Mann-Whitney U Test to compare aCO_2_ versus eCO_2_ using p < 0.01 as threshold for significance and log2FC values |1.5|. Heatmap clustering were performed with these significant masses and were plotted chronologically without hierarchical clustering on the replicates to locate clusters among CO_2_ conditions in each month by utilising RStudio (version 1.4.1103) packages dplyr, viridis and pheatmap. Scripts are available at https://github.com/PlantPriming/MatureOakCO2.

#### 4.5 Metabolites identification

MarVis (Version 6.7.3) suite was used for pathway and compound identification for the significant m/z. Significant differential masses by the Mann-Whitney U Test comparing aCO_2_ versus eCO_2_ per month were selected and combined in a single .csv file containing their peak intensities values, retention time (RT) and m/z. The resulting file was uploaded to the MarVis software (Version 6.7.3). Putative identifications (MS1 level) (Schymanski *et al*., 2014) and pathway enrichment analysis were conducted by MarVis-Pathway using the Kyoto Encyclopedia of Genes and Genomes (KEGG) *Populus trichocarpa* (poplar)-filtered database, and two internal libraries kindly shared by Dr. Victoria Pastor research group for metabolite identification (Gamir *et al*., 2014; Schymanski *et al*., 2014). For the identification thresholds of 0.01 for the m/z difference and 5 s for RT were used. MarVis-Pathway Compound identifications were downloaded and selected based on a p < 0.05 (Kruskal-Wallis) with FDR Benjamin-Hochberg (BH) calculated for the enrichment pathway analysis.

## Results

### Overview of biotic stress incidence across multiple years at the BIFoR- FACE facility

The general incidence of powdery mildew (PM) in the BIFoR FACE facility was relatively stable between 2016 and 2019, thereafter becoming much more variable with a tendency to be lower (Figure 1A). In 2020, disease incidence was markedly lower and represents the lowest disease incidence recorded across the study period (Figure 1A). A statistically significant reduction in PM incidence was also observed in the years from 2021-2024 in comparison to 2016-2019 with each of these years exhibiting disease levels lower than the previous one. Seasonal analysis of disease incidence within each year revealed a distinct pattern with May and September consistently exhibiting lower disease incidence compared to the peak months of the growing season (Figure 1B). Statistically significant differences were observed, with disease incidence being significantly lower in May compared to July and August, and September showing lower levels than August. Out of the three months with high levels (i.e. June, July and August), August was the month with the highest PM incidence. May was the most variable month, likely due to the presence of samples not yet showing disease symptoms (Figure 1B).

**Figure 1.**
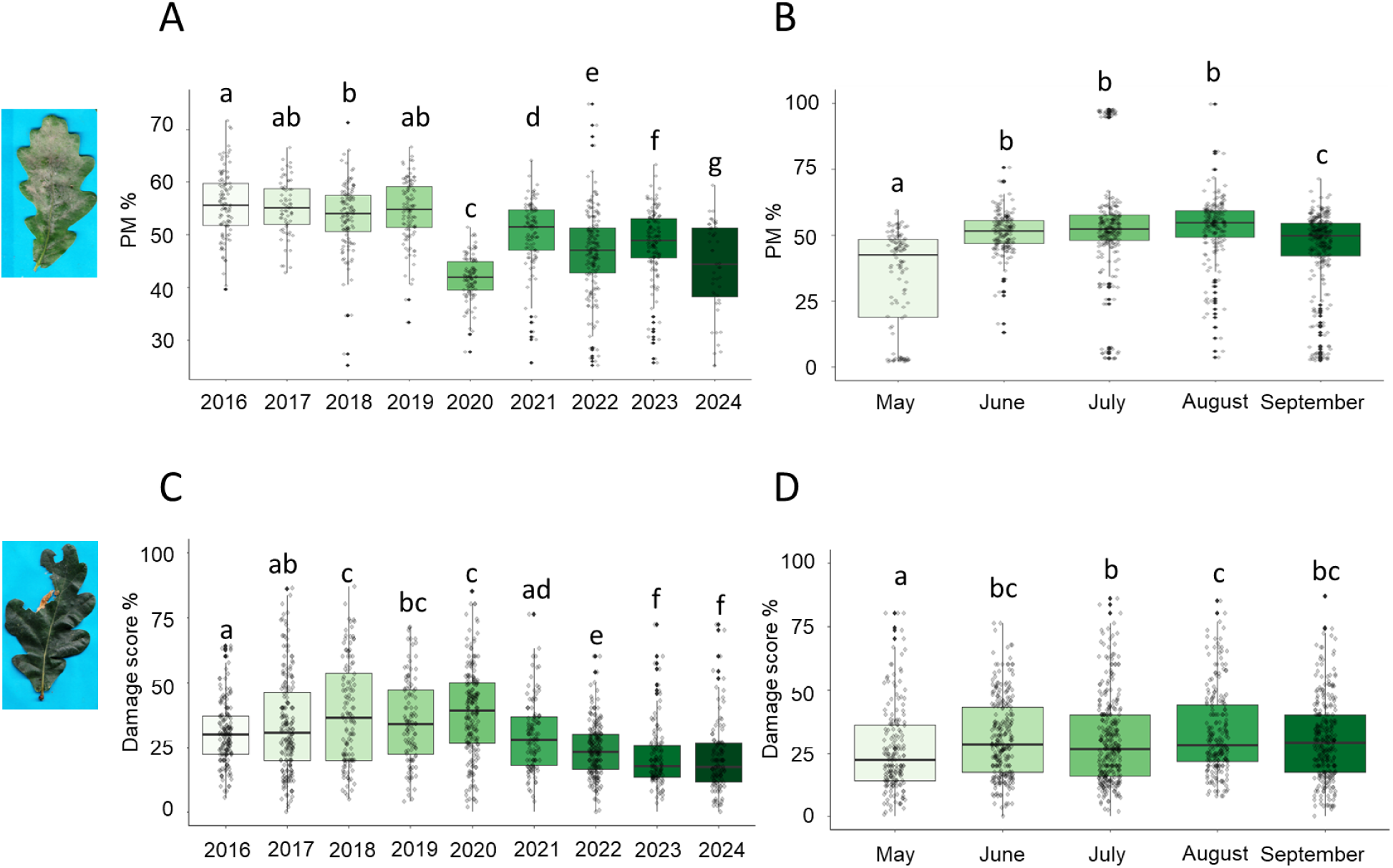
General incidence of powdery mildew (PM) and insect damage across multiple years and within the growing season combining ambient and elevated CO_2_ arrays. A) Yearly incidence of PM expressed as PM percentage (%) from 2016 to 2024. B) Seasonal variation in PM incidence from May to September. C) Yearly incidence of insect damage expressed as % damage score from 2016 to 2024. D) Seasonal variation in insect damage from May to September. Different letters indicate statistically significant differences between years and months in each specific graph (Wilcoxon rank-sum test; p < 0.05 Benjamini-Hochberg (BH) adjusted; n = 115-189). Box plots indicate the 75 and 25 percentiles, middle lines indicate the median and the length of the box is the interquartile range (IQR). Whiskers indicate 1.5x the IQR above and below the mean, and each black circle indicates a data point, considered a biological replicate. Colours in the box plots are used to highlight the progression of time, with darker colours indicating most recent years (A, C) or months throughout the growth season (B, D).

Similarly to PM, while insect damage remained relatively stable from 2016 to 2020, a statistically significant decline was observed from 2021 onward, with each subsequent year i.e. 2022 to 2024 showing a further statistically significant reduction in damage incidence (Figure 1C). The seasonal distribution of insect damage incidence indicated that August had the highest levels of insect damage throughout the study period (Figure 1D). Small but statistically significant differences were observed between months, with August showing significantly higher damage incidence compared to May and September. These findings highlight that while a reduction in biotic stress incidence has been observed over time, contrasting temporal patterns between PM and insect damage incidence suggest different underlying drivers or responses to environmental factors.

### Overall powdery mildew disease incidence under eCO_2_ and aCO_2_ conditions

Analysis of PM incidence across the CO_2_ enrichment period (2017–2024) revealed no significant differences between aCO_2_ and eCO_2_ conditions when considering the dataset as a whole (Figure 2A). Mean PM % scores were nearly identical between the two treatments, with values of 47.64% under aCO_2_ and 47.56% under eCO_2_ (Figure 2A). Similarly, when treating individual trees as biological replicates (i.e. using canopy levels as technical replicates), we obtained values of 49.47% for aCO_2_ and 49.41% in eCO_2_ (Figure 2B), again with no significant differences observed (Figure 2A,B). This trend was consistent across the different experimental arrays, where no major statistically significant differences were detected. Whereas A5 (ambient) showed lower numbers of PM infection in comparison to A1-A4, these differences were not significant against A6 (elevated) (Figure 2C). Further analysis of disease severity at different canopy levels showed no significant impact of eCO_2_, with disease incidence remaining stable across the top, middle and bottom sections of the canopy (Figure 2D-F). Additionally, when average PM% scores were assessed across the years, results reinforced the consistency between aCO_2_ and eCO_2_, with no statistically significant differences emerging at any point (Supplementary Figure 2A).

**Figure 2.**
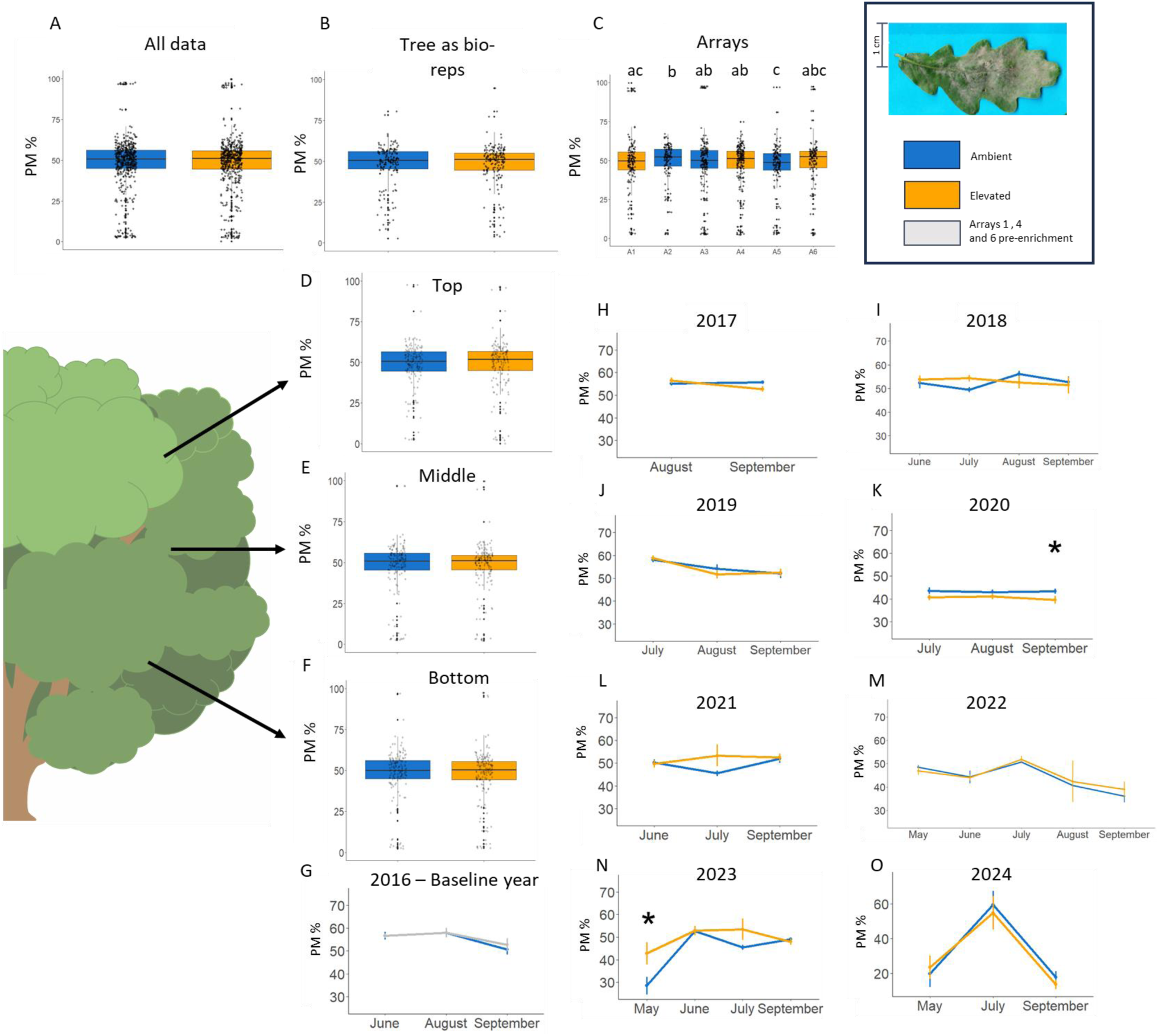
Effect of elevated CO_2_ (eCO_2_) on powdery mildew (PM) incidence at different spatial and temporal scales. A) Overall comparison of PM % between ambient CO_2_ (aCO_2_) and eCO_2_ across all years of CO_2_ enrichment (2017–2024). Grey circles indicate individual data points/images considered as biological replicates (n= 844 for aCO_2_ - 887 for eCO_2_). B) PM incidence analysed with individual trees as biological replicates. Grey circles indicate data per each tree, combining the three canopy levels (n= 295 for aCO_2_ - 296 for eCO_2_). C) PM severity across different experimental arrays 1 to 6 (A1–A6) under aCO_2_ and eCO_2_ conditions (n= 297 for A1, 267 for A2, 312 for A3, 343 for A4, 265 for A5 and 247 for A6). Different letters indicate statistically significant differences between arrays (Wilcoxon rank-sum test; p < 0.05 Benjamini-Hochberg (BH) adjusted). D) PM % scores at the top of the canopy (n= 281 for aCO_2_ - 297 for eCO_2_). E) PM % scores at the middle of the canopy (n= 286 for aCO_2_ - 296 for eCO_2_). F) PM % scores at the bottom of the canopy (n= 282 for aCO_2_ - 294 for eCO_2_). G-O) Year-by-year breakdown of PM % from 2016 (pre-enrichment) through to 2024. Asterisks indicate statistically significant differences between eCO_2_ and aCO_2_ in the years in each graph (Kruskal-Wallis; p < 0.05; n= 844-887). Box plots indicate the 75 and 25 percentiles, middle lines indicate the median and the length of the box is the interquartile range (IQR). Whiskers indicate 1.5x the IQR above and below the mean, and each black circle indicates a data point, considered a biological replicate.

Year-specific analyses confirmed that, prior to the start of CO_2_ enrichment in 2016, PM incidence was similar across all arrays (Figure 2G). Following the initiation of eCO_2_ in 2016, PM incidence remained comparable between aCO_2_ and eCO_2_ groups for most years (Figure 2H-J, L, M, O), with two key exceptions. In 2020, disease severity was significantly lower under eCO_2_ (39.98%) compared to aCO_2_ (42.47%) (p = 0.009) across all months (Figure 2K). Conversely, in May 2023, an opposite trend was observed, with PM incidence being significantly higher under eCO_2_ (42.74%) than aCO_2_ (28.51%) (p = 0.002) (Figure 2N). However, no further differences were found in the months assessed in 2023 after May.

Overall, these findings indicate that eCO_2_ does not exert a consistent effect on PM incidence, as PM% remained stable across treatment groups, years, canopy levels, and experimental arrays. However, isolated instances of significant differences in disease incidence suggest potential interactions between eCO_2_ and environmental conditions in specific years.

### Overall insect damage under eCO_2_ and aCO_2_ conditions

The long-term assessment of insect damage from 2017 to 2024 under eCO_2_ conditions revealed no significant deviation from the aCO_2_ treatment when considering the dataset as a whole (Figure 3A). The mean percentage of insect damage remained nearly identical between treatments, with aCO_2_ trees showing an average of 28.68% damage and eCO_2_ trees exhibiting 30.35% (Figure 3A) however, these differences were not statistically significant. Likewise, when individual trees were analysed as biological replicates, no statistically significant differences emerged between the two CO_2_ conditions (Figure 3B), further emphasising the similarity between conditions. An exception to this pattern was detected when insect damage was compared across the experimental arrays (A1–A6). Array 6 (A6; eCO_2_) exhibited lower insect damage scores compared to the other arrays apart from the neighbouring A5 (aCO_2_), suggesting that localised environmental factors or tree-specific characteristics may be influencing herbivory rates at that spatial scale (Figure 3C). Despite this variation among arrays, damage incidence remained uniform across different canopy positions (top, middle, and bottom), with no significant treatment-related differences observed (Figure 3D–F). Effect size analysis further confirmed that the influence of eCO_2_ on herbivory was negligible at all canopy levels, reinforcing the broader finding that insect feeding patterns were largely unaffected by eCO_2_ conditions.

**Figure 3.**
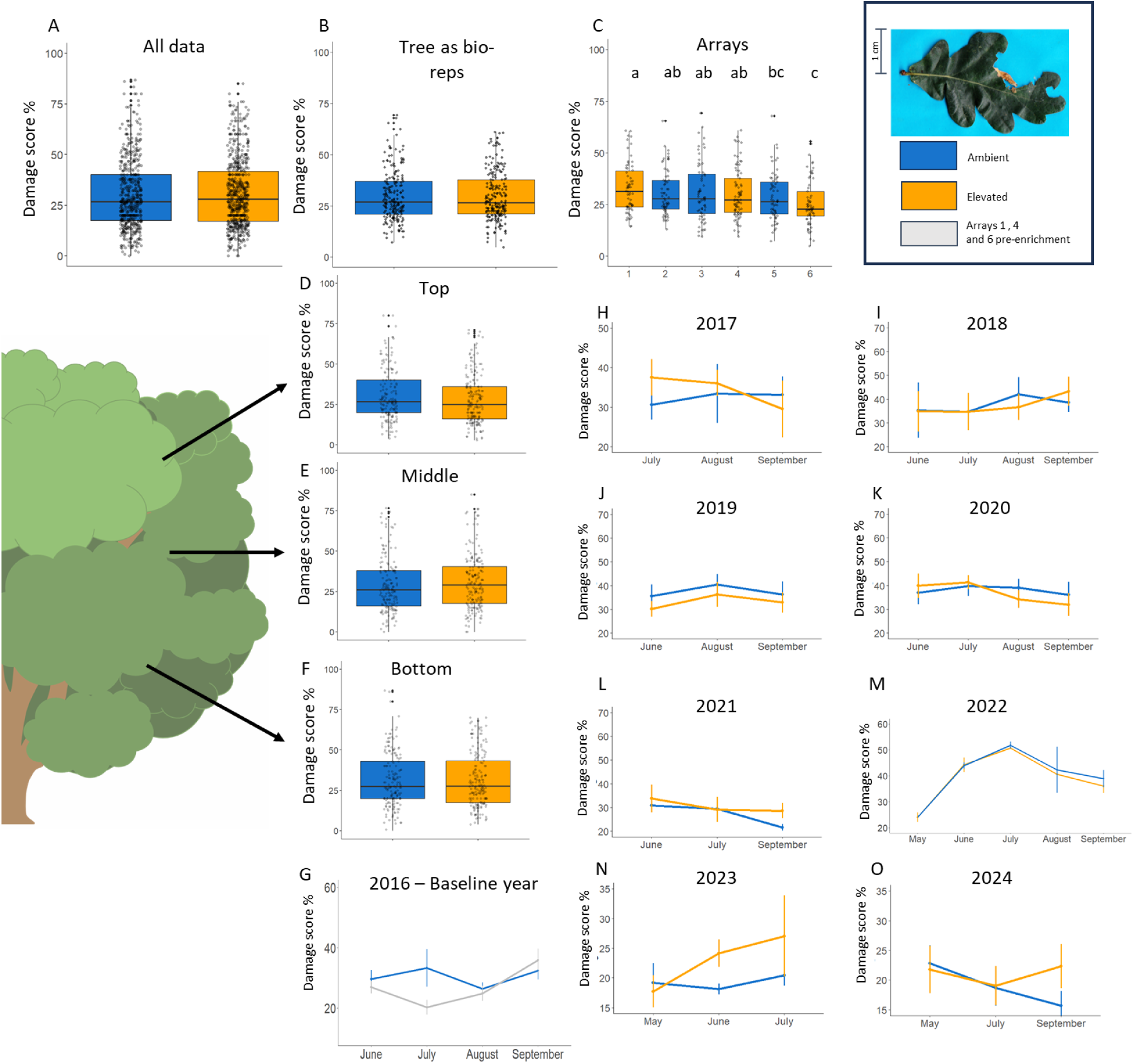
Effect of elevated CO_2_ (eCO_2_) on insect damage scores at different spatial and temporal scales. A) Overall comparison of insect damage between ambient CO_2_ (aCO_2_) and eCO_2_ across all years of CO_2_ enrichment (2017–2024). Grey circles indicate individual data points/images considered as biological replicates (n=657 for aCO_2_ - 706 for eCO_2_). B) Insect damage analysed with individual trees as biological replicates. Grey circles indicate data per each tree, combining the three canopy levels (n= 219 for aCO_2_ - 235 for eCO_2_). C) Insect damage severity across different experimental arrays (A1–A6) under aCO_2_ and eCO_2_ conditions (n= 219 for A1, 228 for A2, 211 for A3, 270 for A4, 210 for A5 and 225 for A6). Different letters indicate statistically significant differences between arrays (Wilcoxon rank-sum test; p < 0.05 Benjamini-Hochberg (BH) adjusted). D) Insect damage at the top of the canopy (n= 218 for aCO_2_ - 236 for eCO_2_). E) Insect damage at the middle of the canopy (n= 217 for aCO_2_ - 236 for eCO_2_). F) Insect damage at the bottom of the canopy (n= 218 for aCO_2_ - 226 for eCO_2_). G-O) Year-by-year breakdown of insect damage from 2016 (pre-enrichment) through to 2024. Box plots indicate the 75 and 25 percentiles, middle lines indicate the median and the length of the box is the interquartile range (IQR). Whiskers indicate 1.5x the IQR above and below the mean, and each black circle indicates a data point, considered a biological replicate.

Examining damage trends over time revealed no divergence between the eCO_2_ and aCO_2_ treatments. When average annual insect damage scores were assessed (Supplementary Figure 2B), the two groups followed nearly identical trajectories, with no statistically significant differences emerging across years. Similarly, baseline insect damage levels in 2016, prior to CO_2_ enrichment, showed occasional differences but no statistically significant variation between treatment groups (Figure 3G). From 2017 onward, some fluctuation in damage severity was observed across years, but no clear or sustained effect of eCO_2_ was detected (Figure 3H–O). These results suggest that while spatial variability in herbivory may exist within the study site, particularly in specific arrays such as A6, eCO_2_ does not appear to drive changes in overall insect damage patterns over time.

### Biochemical analysis of photosynthetic metabolites

To test whether eCO_2_ influences the accumulation of photosynthetic metabolites in ways that could affect defence responses, we conducted a targeted analysis of key pigments and antioxidant compounds. No statistically significant differences in mean abundance of photosynthetic metabolites were observed between aCO_2_ and eCO_2_ treatments based on post-hoc Tukey tests (Figure 4). However, differences in the variability of metabolite levels between treatments were evident. In general, eCO_2_ trees exhibited greater variability across nearly all metabolites analysed. This pattern was particularly pronounced for glutathione (Figure 4A) and cysteine (Figure 4B), where replicate values in the eCO_2_ group spanned a broader range compared to the more uniform distributions observed under aCO_2_. Other compounds (Figure 4C- J) such as lutein (Figure 4C), neoxanthin (Figure 4D), chlorophyll a (Figure 4E), chlorophyll b (Figure 4F), and b-carotene (Figure 4G) showed all the same patterns with different levels of variability. Interestingly, antheraxanthin (Figure 4K) and a-carotene (Figure 4L) were the exception to this trend, with slightly reduced variability under eCO_2_ conditions. This widespread increase in variability under eCO_2_ suggests a heterogeneous physiological response among trees, potentially reflecting individual-level plasticity in adjusting pigment and antioxidant pools under altered carbon availability.

**Figure 4.**
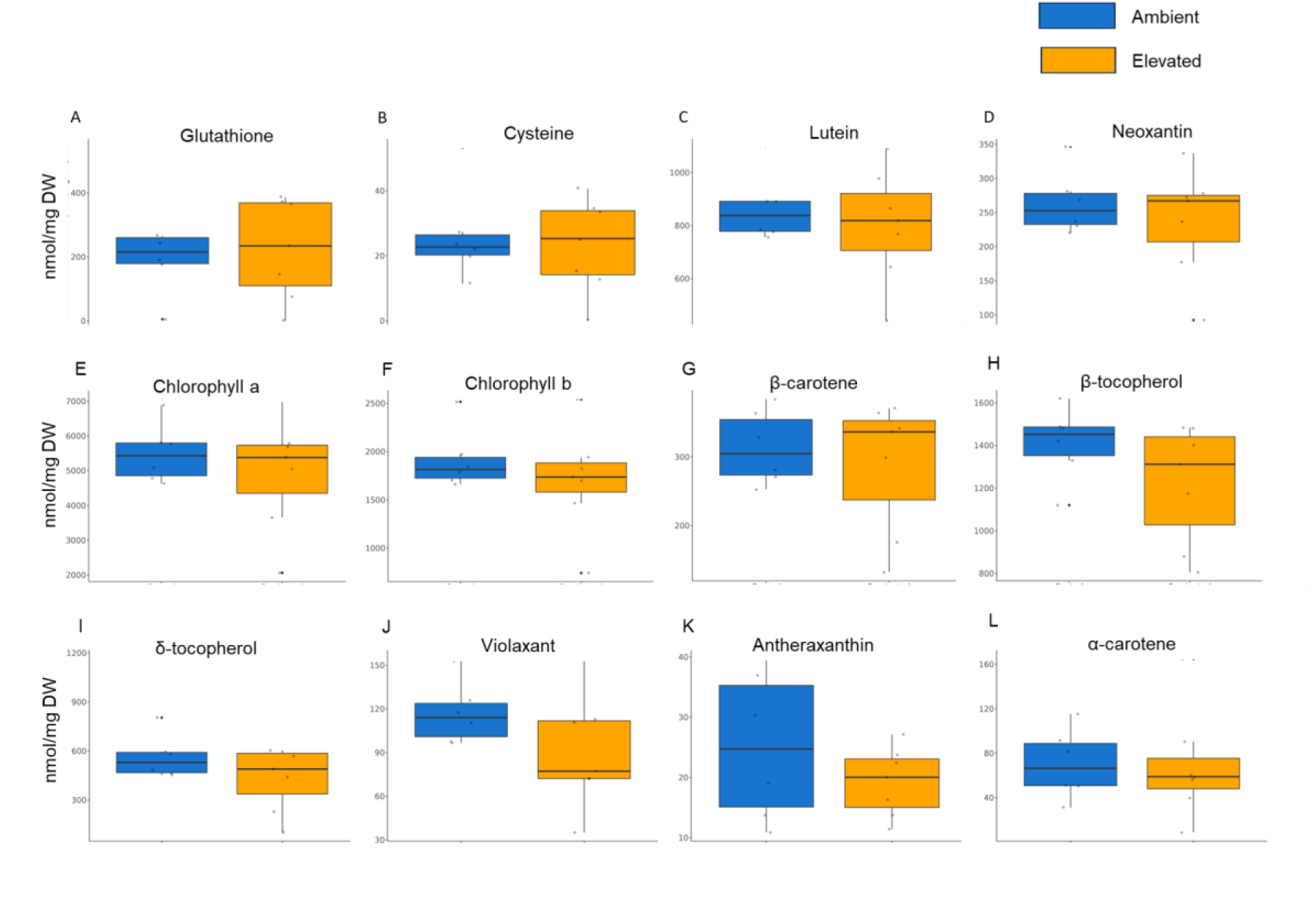
Targeted analysis of key photosynthetic pigments and antioxidant metabolites in oak leaves collected under ambient CO_2_ (aCO_2_) and elevated CO_2_ (eCO_2_) conditions. Each panel shows individual replicate values (n = 6) and group means for: A) Glutathione, B) Cysteine, C) Lutein, D) Neoxanthin, E) Chlorophyll a, F) Chlorophyll b, G) β-carotene, H) α-tocopherol, I) d-tocopherol, J) Violaxanthin, K) Antheraxanthin, and L) α-carotene. Box plots indicate the 75 and 25 percentiles, middle lines indicate the median and the length of the box is the interquartile range (IQR). Whiskers indicate 1.5x the IQR above and below the mean. Not statistically significant data found (Tukey test; p < 0.05; n = 6).

### Temporal changes in the oak metabolome under elevated CO_2_

To test the effect of eCO_2_ in the metabolomic profiles of the trees, leaf samples were collected from May to September of 2021 and subjected to untargeted metabolomic analysis. Considering that no differences in canopy levels were identified in PM and insect incidence, only mid-canopy samples were analysed. A total of 12,299 and 10,446 m/z features were detected in positive and negative ionisation modes, respectively. To investigate differences between aCO_2_ and eCO_2_, PCA plots were created in each collection month using the combined (ESI- and ESI+) dataset. PCA analyses revealed very few differences across the sampling season with the exception of September, where some differences were identified (Figure 5A). In May, 60 m/z features were differentially accumulated, rising to 142 in June and 153 in July, and peaking in September with 279 features (Figure 5B). This trend suggests that the impact of eCO_2_ on the oak metabolome intensifies as the season progresses, likely reflecting the cumulative physiological effects of prolonged CO_2_ exposure over time. However, this rising trend was not observed in August, where there was a notable drop, with only 68 features observed (Figure 5B).

**Figure 5.**
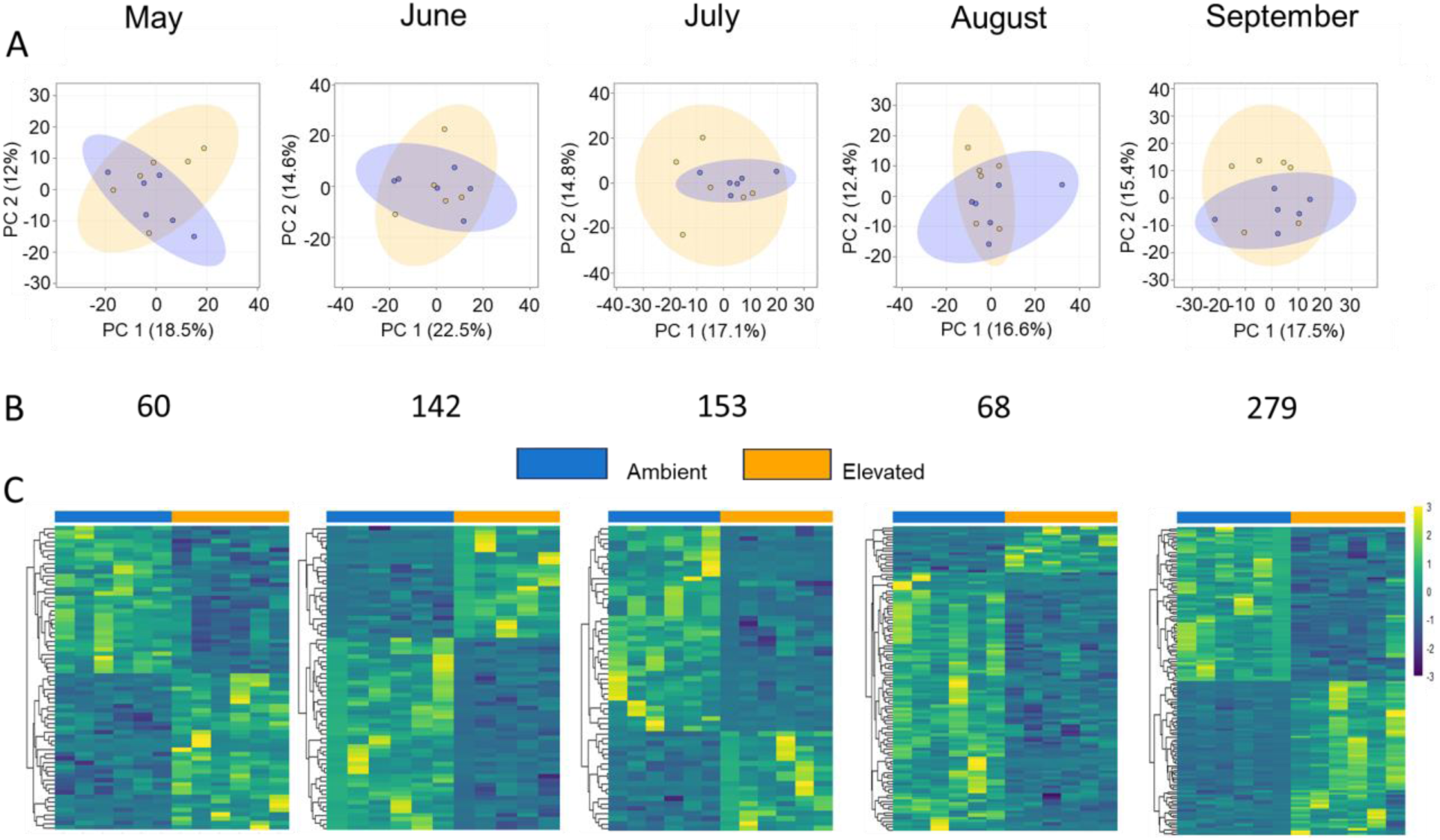
Temporal patterns of differentially accumulated m/z features between ambient CO_2_ (aCO_2_) and elevated CO_2_ (eCO_2_) trees across the 2021 growing season. A) Principal component analysis (PCA) of untargeted metabolomic profiles across monthly collections from May to September. Each panel represents a separate month, with aCO_2_ and eCO_2_ samples shown. B) Number of differentially accumulated mass-to-charge (m/z) features between aCO_2_ and eCO_2_ per month (p < 0.01 and log2FC |1.5|). C) Heatmaps showing the relative abundance of differentially accumulated m/z values for each month.

A combined PCA containing the data from all studied months further supported these temporal dynamics (Supplementary Figure 3A). In May, both aCO_2_ and eCO_2_ samples separated clearly from other months along PC1 (18.8%), indicating early-season metabolic distinctiveness. Meanwhile, PC2 (5.4%) appeared to reflect a seasonal progression, potentially linked to increasing biotic stress such as pathogen or insect pressures. Notably, June samples under eCO_2_ exhibited a high degree of variability, with data points widely dispersed across PCA space. This variability diminished over the following months, with August samples clustering tightly.

To explore the structure of metabolomic differences further, clustering analyses were conducted using the differentially accumulated m/z values per month. Heatmap representations of monthly profiles revealed a consistent trend of downregulation under eCO_2_: 32 downregulated, 28 upregulated in May; 83 down, 59 up in June; 104 down, 49 up in July; and particularly pronounced in August with 51 down, 17 up (Figure 5C). However, this pattern shifted in September, where a higher proportion of m/z features were upregulated in eCO_2_ samples, with 112 down, 167 up. When clustering was performed using all months, two dominant clusters emerged, largely driven by the distinct metabolic signature of May (Supplementary Figure 3B). To disentangle the seasonal from the treatment effects, May was excluded and samples from both CO_2_ conditions were combined. This analysis showed that clustering was determined primarily by month rather than CO_2_ treatment (Supplementary Figure 3C), underscoring the strong seasonal influence on the oak metabolome. Finally, analysis of the m/z values that changed with eCO_2_ across all the months of June to September, unravelled that no m/z features were shared throughout the season (Supplementary Figure 3D), therefore indicating that there is not a common signature of eCO_2_ effects in the metabolome of mature oaks. Taken together, these results indicate that while eCO_2_ induces measurable changes in metabolite accumulation, particularly in September, these effects are not consistent across the growing season. Instead, temporal dynamics, rather than CO_2_ treatment alone, appear to be the dominant driver of variation in the oak metabolomic profile.

A total of 691 differentially accumulated m/z features in response to the eCO_2_ along May-September (399 ESI+ and 292 ESI-) were selected to perform the identification and enrichment pathway analysis. A total of 66 metabolites were identified at the MS1 level (Supplementary Table 2), which represents around 10% of the selected dataset. From the 66 compounds identified, an overrepresentation of amino acids and derivatives was observed (13 out of 66) as well as alkaloids (6 out of 66) and phenylpropanoids (7 out of 66). Moreover, identification allocated different compounds in the glucosinolate metabolism (9 out of 66). Out of those 9, 2 were also annotated into the 2-oxocarboxylic acid metabolism pathway. Considering that oak trees do not have glucosinolates, the 7 metabolites only allocated to this pathway were not taken into further consideration. The 59 identified compounds were then used for pathway enrichment analysis, which revealed two significantly enriched pathways with FDR p < 0.05: 2-oxocarboxylic acid metabolism (p = 0.01081) and Pantothenate and CoA synthesis (p= 0.0297) (Supplementary Table 2).

## Discussion

Understanding how elevated atmospheric CO_2_ (eCO_2_) influences forest health and resilience is a pressing priority in the face of accelerating climate change. This study represents one of the most comprehensive assessments to date of the long-term effects of eCO_2_ on mature oak (*Quercus robur*) trees, combining nine years of biotic stress monitoring with both targeted and untargeted metabolomic profiling in a fully operational temperate forest ecosystem. We hypothesised that extended exposure to eCO_2_ would alter susceptibility to powdery mildew (PM) and insect herbivory, and induce distinct metabolic adjustments, particularly within photosynthetic and antioxidant pathways. While we observed subtle, temporally dynamic biochemical shifts, our findings demonstrate that eCO_2_ did not significantly alter biotic stress across the duration of the study.

Before assessing the impact of eCO_2_ on disease incidence, we first aimed to understand the baseline patterns of powdery mildew (PM) and insect herbivory in the FACE facility over the course of the study (2016–2024). Multi-year monitoring revealed a clear downward trend in both PM incidence and herbivory damage, particularly from 2020 and 2021 onward, respectively.

Notably, 2018 stood out for its unusually high variability in insect damage, coinciding with a mass defoliation event caused by the late hatch of oak winter moth caterpillars, triggered by the "Beast from the East" cold snap. This event highlights how climatic anomalies can disrupt longer-term biotic stress patterns, emphasizing the need to account for extreme events in climate-forest interaction studies. Further spatial analysis within the FACE facility also revealed consistent differences among experimental arrays, with A5 and A6 showing lower levels of PM and insect damage compared to other areas (Figures 2C and 3C). Given that these arrays are situated close together within the Mill Haft Estate (Hart *et al*., 2020), local microclimatic differences such as variations in light availability, humidity, and wind exposure likely contribute to these spatial patterns. Seasonal analysis highlighted that August consistently showed the highest PM incidence. This peak is particularly interesting given that August typically coincides with low soil moisture levels and reduced leaf water availability, conditions that could favour PM infection dynamics. PM development is known to thrive under moderate temperatures (15–25°C) and high relative humidity (78–90%) (Pap *et al*., 2013), while heavy rainfall can negatively affect PM by washing spores off leaves. August conditions at the FACE site often match these optimal ranges for PM development, particularly with intermittent dry spells that reduce spore wash-off but maintain high humidity within the dense canopy. Thus, the environmental conditions typical of late summer may significantly contribute to the higher PM pressure observed during this period. Taken together, these findings suggest that while biotic stress in mature oaks has generally decreased over time, PM and insect damage are strongly shaped by seasonal, interannual, and spatial variability, with potential sensitivity to specific environmental triggers such as extreme weather events, local microclimate, and seasonal drought patterns.

Throughout the enrichment period (2017–2024), no sustained differences were detected between eCO_2_ and ambient CO_2_ (aCO_2_) trees in PM or insect damage. Isolated events such as lower PM incidence in eCO_2_ trees in 2020 and higher incidence in May 2023 were not sustained across the growing season or reproduced in other years (Figure 2). Intriguingly, 2020 was the year of overall lowest PM pressure (Figure 1A), and we speculate that under low disease pressure, trees growing in eCO_2_ may allocate surplus carbon to increase defence responses, potentially through mechanisms similar to priming. Previous research has shown that disease pressure is a key determinant of defence activation, specifically when it comes to priming and the activation of SA- and JA-dependent pathways (Pétriacq *et al*., 2016). Moreover, eCO_2_ has been implicated in enhancing priming against biotrophic pathogens (Williams *et al*., 2018). Therefore, we could speculate that eCO_2_ activates priming of defences, which are capable of enhancing disease resistance when the disease pressure is low. Conversely, in May 2023, when disease symptoms were just beginning to emerge and overall pressure was also low, we observed a reversed phenotype, with eCO_2_ trees displaying higher PM incidence. It is plausible that early in the season, trees allocate more resources toward growth (Giolai & Laine, 2024), delaying investment in defence and thereby increasing susceptibility under eCO_2_. Importantly, PM was not detected in May 2020, and June material was not available, making it difficult to draw firm conclusions about seasonal disease pressure. Overall, these findings suggest that such anomalies likely reflect interactions between environmental conditions and tree physiology, rather than consistent treatment effects.

Insect herbivory mirrored these findings. Damage levels remained stable across treatments, with no significant impact of eCO_2_ (Figure 3). Damage patterns were also consistent across canopy layers, suggesting that vertical leaf position does not influence stress susceptibility under eCO_2_. These results contrast with prior studies in herbaceous or juvenile woody plants, where eCO_2_ has been linked to altered pathogen and herbivore interactions (Pugliese *et al*., 2010; Kazan, 2018) but is consistent with Roberts *et al*. (2022) who found no significant change in herbivory during the first three years of eCO_2_ at the FACE facility. It is important to note that most eCO_2_ studies typically involve highly controlled lab conditions, using young plants, and unrealistically high CO_2_ concentrations. Our findings underscore the importance of studying mature trees in complex field environments, where buffering capacity and ecological context likely moderate CO_2_ effects.

While biotic stress responses remained relatively stable across treatments, metabolomic profiling revealed important physiological differences, particularly in compound variability. In our targeted analysis, no significant differences in mean metabolite abundance were detected between aCO_2_ and eCO_2_ conditions. However, greater variability among eCO_2_ samples was observed for several key metabolites involved in photosynthesis and redox regulation, including glutathione, cysteine, chlorophylls, lutein, and neoxanthin (Figure 4). This increased variability likely reflects individual-level physiological plasticity in response to eCO_2_. This is in contrast with a recent study at the FACE facility, where eCO_2_ was found to increase chlorophyll content (Gardner *et al*., 2022). In this study over 5000 *in situ* chlorophyll absorbance measurements were performed, therefore providing a much deeper understanding on the effect of eCO_2_ in these photosynthetic pigments. Interestingly, not all compounds followed the high variation pattern under the eCO_2_ treatment. β- carotene and antheraxanthin, both part of the xanthophyll cycle, showed stable levels across treatments, suggesting that these pigments may be more tightly regulated, potentially due to their critical roles in photoprotection and oxidative stress mitigation (Fernandez-Marin *et al*., 2021). The distinct behaviour of these metabolites highlights the possibility of compound-specific regulation pathways depending on their functional roles in leaf physiology. Moreover, it is notable that samples for this analysis were collected in August, the month with the highest observed levels of both PM and insect damage (Figure 1B, D). Previous studies have shown that PM infection in oak can elevate ascorbate-glutathione cycle enzyme activity, phenolic compounds, and metal accumulation (Skwarek-Fadecka *et al*., 2024), all associated with defensive responses. Therefore, we speculate that the higher variability of redox-related metabolites under eCO_2_ may be linked to differential stress responses among trees experiencing varying degrees of biotic pressure. In contrast, the stability of certain xanthophylls could reflect their primary role in responding to light stress, rather than pathogen or herbivore attack. Finally, these metabolite-level differences may also be influenced by leaf developmental stage or microenvironmental variation, both of which can modulate metabolic outputs even within a single canopy.

Untargeted metabolomics provided additional insight. The number of differentially accumulated metabolites increased throughout the season, peaking in September, pointing to a cumulative effect of long-term eCO_2_ exposure. Interestingly, this upregulation pattern was not observed in August, possibly reflecting a unique physiological state at that point in the season or the biotic stress pressure reducing the effects of eCO_2_ exposure. Principal component analysis also highlighted that May samples were metabolically distinct, and eCO_2_ samples in June showed high variability, which decreased later in the season. This suggests a seasonally gated or acclimatory metabolic response, where trees adjust to eCO_2_ progressively across the season. Untargeted metabolomic analysis provided additional insight into the temporal dynamics of eCO_2_ responses. The number of differentially accumulated metabolites increased progressively over the growing season, peaking in September, suggesting a cumulative effect of long-term eCO_2_ exposure. However, this trend did not hold in August, when the number of affected metabolites dropped markedly. This divergence may reflect a unique physiological state during this period or a shift in metabolic priorities due to elevated biotic stress pressure, which was highest in August (Figure 1B,D). Principal component analysis supported these seasonal patterns. May samples appeared metabolically distinct, while eCO_2_ samples in June exhibited high within-group variability, which gradually diminished over the season. This trend points to a possible seasonally gated or acclimatory metabolic response, wherein trees adjust their biochemical profile to eCO_2_ progressively over time. An additional explanation for the August anomaly may lie in sampling-related factors. Oaks are known to produce secondary leaf flushes, such as lamma growth, particularly in late summer (McKown *et al*., 2013). It is possible that younger, newly emerged leaves were collected in August, differing developmentally from earlier samples and potentially altering metabolomic profiles. This may also account for the discrepancies observed between targeted and untargeted analyses during this month. Finally, it is plausible that in August, when biotic stress pressure peaks, oak trees redirect metabolic resources toward defence, thus reducing the observable impact of eCO_2_ on primary metabolism (Supplementary Table 2). Future work should aim to differentiate between primary and secondary leaf cohorts, and to explore how leaf age and developmental stage influence the metabolomic response to eCO_2_ and concurrent environmental stressors.

Despite a relatively low number of annotated m/z features (Supplementary Table 2), monthly metabolite profiling revealed that eCO_2_ induces broad alterations across the oak leaf metabolome. These changes affected both primary and secondary metabolic pathways, indicating a wide-ranging physiological response. Primary metabolites impacted by eCO_2_ included amino acids, CoA derivatives, folate-related compounds, and intermediates of the 2-oxocarboxylic acid cycle, molecules central to energy metabolism, cellular function, and biosynthetic activity (Smith *et al*., 2007; Maurino & Engqvist, 2015). These changes may be linked to enhanced photosynthetic rates typically observed under eCO_2_ in oak trees. Seasonal accumulation patterns of amino acid-related metabolites, especially during June and July, aligned with the peak of photosynthetic activity (Norby *et al*., 2024), suggesting increased metabolic flux to support protein synthesis, growth, and stress responses. Enhanced soil N mineralization and plant available N to trees under eCO_2_ may have supported the flexibility in leaf amino acids and N bearing metabolomes (Sgouridis *et al*., 2023; Rumeau *et al*., 2024). Many of these compounds may also act as precursors to phenylpropanoids and other defence-associated secondary metabolites (El-Azaz *et al*., 2023).

Together, this reinforces the interpretation that eCO_2_ primarily modulates core metabolic processes, especially during active growth phases.

In parallel, secondary metabolites, including alkaloids, flavonoids, phenylpropanoids, and terpenoids, showed seasonally dynamic patterns. For instance, shifts in alkaloid profiles, such as increased anti-digestive and microbial-deterrent compounds (Özer *et al*., 2017), suggest a reprogramming of defence metabolism in response to environmental cues, including biotic stress. These trade-offs may reflect changes in defence priorities across the season. Several redox-active metabolites, including glutathione precursors and sulphur-containing intermediates, were also responsive to eCO_2_. Their variability may reflect tree-specific plasticity in oxidative stress regulation and immune priming. Interestingly, a general trend of downregulation was observed for some metabolites, including those involved in suberin and wax biosynthesis, key components of the plant’s physical barrier against pests and pathogens (Serrano *et al*., 2014). While these changes could theoretically impact resistance, we did not observe any consistent differences in disease or insect susceptibility under eCO_2_ (Figure 2, Figure 3). This suggests that the metabolic shifts identified here, though biologically relevant, may not be sufficient to elicit a phenotypic change in biotic stress resistance, at least under the prevailing environmental conditions.

Overall, our untargeted metabolome analysis indicates that eCO_2_ subtly reshapes the metabolic landscape of mature oak trees, with a greater influence on primary metabolism than on specialised secondary pathways. These shifts appear to support a flexible strategy that balances growth and maintenance during periods of high photosynthetic activity, while also allowing for metabolic reallocation in response to seasonal biotic and environmental pressures.

In summary, our findings provide a critical counterpoint to previous research suggesting that eCO_2_ disrupts plant defence. In mature oak trees under real-world field conditions, eCO_2_ enrichment of +150 ppm did not increase susceptibility to pathogens or herbivores, nor did it lead to consistent alterations in photosynthetic or antioxidant pathways. Instead, our results suggest a remarkable resilience in mature oaks to moderate eCO_2_, with only subtle, temporally dependent metabolic adjustments. However, the lack of consistent treatment effects does not imply eCO_2_ is inconsequential. Rather, it emphasises that tree responses are highly context-dependent, shaped by interacting climatic, biological, and edaphic variables.

Crucially, ongoing research at BIFoR-FACE is now extending into long-term, multifactorial experiments, incorporating drought, soil biology and biotic challenges alongside CO_2_ enrichment. These efforts will be vital to reveal how mature trees and broader forest systems respond to the complex, overlapping stressors of climate change. Future work should continue to explore multi-stressor dynamics, assess long-term physiological and reproductive consequences and investigate the responses of understorey vegetation and juvenile cohorts, which may differ in sensitivity and resilience compared to mature canopy trees. Together, these insights will be essential for developing informed, adaptive strategies for forest conservation and management in a changing atmosphere.

## Acknowledgments

We thank the wider research community of BIFoR for the extremely valuable discussions on this work. Moreover, we are extremely grateful to the technical team of BIFoR-FACE for their helpful assistance. Specially, we would like to thank Gael Denny for her help and assistance with the monthly green leaf collections. In addition, we thank all the BIFoR volunteers that have helped throughout the years with the monthly green leave collections. Finally, we thank the SCIC of the Universitat Jaume I for technical assistance in metabolomics. This work was funded by the JABBS foundation grant “Resistance strategies of oak trees in the arms race with pathogens“ to EL and RSL and the UKRI grant “MEMBRA” NE/V021346/1 to EL, ARM and SALH. The work of MR was supported by the leverhulme PhD training network Forest Edge. The work of VP was supported by the Valencian grant CIACO/2021/092. SALH is supported by a Leverhulme Research Fellowship RF-2024-396/2. ARM and CM were additionally supported by the UK Natural Environmental Research Council through grant NE/S015833/1 (QUINTUS). FACE funding was provided by the JABBS foundation, the University of Birmingham, and the John Horseman Trust, and the Ecological Continuity Trust.

## Conflicts of Interest

The authors declare no conflicts of interest.

## Data Availability Statement

The data that support the findings of this study are openly available. All data are available in the different repositories and platforms as follows: R packages and scripts can be found in the public GitHub PlantPriming folder MatureOakCO_2_: https://github.com/PlantPriming/MatureOakCO2.

Metabolomic data have been deposited in MetaboLights (Yurekten et al. 2024) under the identifier XXX (available upon acceptance).

## Author contribution

**RSL** conceived and designed the experimental approach, performed or was involved in all experiments, co-supervised MR, AD, KHawkins, and EAP, coordinated summer students and volunteers, carried out data analysis of the metabolome experiment, and wrote the manuscript; **MR** contributed to the initial manuscript draft, performed or was involved in all experiments, and co-analysed metabolomic data; **AD** contributed to manuscript writing, performed full insect damage scoring, analysed PM and insect data, and produced the figures of the manuscript; **KHawkins** scanned leaves collected between 2016–2021 and solely developed the ImageJ macro for PM scoring; **EAP** performed leaf scanning for samples collected between 2016–2021; **SU** provided intellectual input throughout the study; **ARM** provided intellectual input, set up of the FACE facility, collect baseline samples and early samples, and technical assistance with archived material; **KHart** provided technical support and oversight at the BIFoR-FACE facility; **DB** provided technical assistance with archived material, contributed to leaf scanning and coordinated volunteers; **CM** provided technical assistance with archived material and contributed to leaf scanning; **MS** and **IK** performed the targeted biochemical analysis of photosynthetic metabolites; IK additionally provided intellectual input; **SALH** provided intellectual input and co-supervised MR; **VP** performed untargeted metabolomic analysis; **EL** conceived the study, secured funding, supervised the project, performed experiments, mentored the research team, and contributed to manuscript writing. All authors provided input in the submitted draft of this manuscript.

## Notes

### Competing Interest Statement

The authors have declared no competing interest.

## References

1. Alalouni U, Brandl R, Auge H, Schädler M. 2014. Does insect herbivory on oak depend on the diversity of tree stands? Basic and Applied Ecology 15(8): 685–692. 10.1016/j.baae.2014.08.013

2. Bailly C, Kranner I 2011. Analyses of Reactive Oxygen Species and Antioxidants in Relation to Seed Longevity and Germination. In: Kermode AR ed. Seed Dormancy: Methods and Protocols. Totowa, NJ: Humana Press, 343–367.

3. Baker HS, Millar RJ, Karoly DJ, Beyerle U, Guillod BP, Mitchell D, Shiogama H, Sparrow S, Woollings T, Allen MR. 2018. Higher CO_2_ concentrations increase extreme event risk in a 1.5 °C world. Nature Climate Change 8(7): 604–608. 10.1038/s41558-018-0190-1

4. Braga M, P.M A, Marabesi M, Godoy J. 2006. Effects of elevated CO_2_ on the phytoalexin production of two soybean cultivars differing in the resistance to stem canker disease. Environmental and Experimental Botany 58: 85–92. 10.1016/j.envexpbot.2005.06.018

5. Buchner O, Roach T, Gertzen J, Schenk S, Karadar M, Stöggl W, Miller R, Bertel C, Neuner G, Kranner I. 2017. Drought affects the heat-hardening capacity of alpine plants as indicated by changes in xanthophyll cycle pigments, singlet oxygen scavenging, α-tocopherol and plant hormones. Environmental and Experimental Botany 133: 159–175. 10.1016/j.envexpbot.2016.10.010

6. Crowley LM. 2021. Are insects key drivers of change in woodland systems under climate change? Ph.D., University of Birmingham.

7. Díaz S, Settele, J., Brondízio, E. S., Ngo, H. T., Guèze, M., Agard, J., Arneth, A., Balvanera, P., Brauman, K. A., Butchart, S. H. M., Chan, K. M. A., Garibaldi, L. A., Ichii, K., Liu, J., Subramanian, S. M., Midgley, G. F., Miloslavich, P., Molnár, Z., Obura, D., Pfaff, A., Polasky, S., Purvis, A., Razzaque, J., Reyers, B., Roy Chowdhury, R., Shin, Y. J., Visseren-Hamakers, I. J., Willis, K. J., & Zayas, C. N. (Eds.). 2019. Summary for policymakers of the global assessment report on biodiversity and ecosystem services of the Intergovernmental Science-Policy Platform on Biodiversity and Ecosystem Services. Bonn, Germany: IPBES Secretariat.

8. Dong J, Gruda N, Lam SK, Li X, Duan Z. 2018. Effects of Elevated CO_2_ on Nutritional Quality of Vegetables: A Review. Frontiers in Plant Science Volume 9 - 2018. 10.3389/fpls.2018.00924

9. El-Azaz J, Moore B, Takeda-Kimura Y, Yokoyama R, Wijesingha Ahchige M, Chen X, Schneider M, Maeda HA. 2023. Coordinated regulation of the entry and exit steps of aromatic amino acid biosynthesis supports the dual lignin pathway in grasses. Nat Commun 14(1): 7242. 10.1038/s41467-023-42587-7

10. Erb MM, Stefan; Howe, Gregg A. 2012. Role of phytohormones in insect-specific plant reactions. Trends in Plant Science 17(5). 10.1016/j.tplants.2012.01.003

11. Fernandez-Marin B, Roach T, Verhoeven A, Garcia-Plazaola JI. 2021. Shedding light on the dark side of xanthophyll cycles. New Phytol 230(4): 1336–1344. 10.1111/nph.17191

12. Friedlingstein P, O’Sullivan M, Jones MW, Andrew RM, Hauck J, Landschützer P, Le Quéré C, Li H, Luijkx IT, Olsen A, et al. 2025. Global Carbon Budget 2024. *Earth Syst*. Sci. Data 17(3): 965–1039. 10.5194/essd-17-965-2025

13. Gamir J, Sánchez-Bel P, Flors V. 2014. Molecular and physiological stages of priming: how plants prepare for environmental challenges. Plant Cell Reports 33(12): 1935–1949. 10.1007/s00299-014-1665-9

14. Gardner A, Ellsworth DS, Pritchard J, MacKenzie AR. 2022. Are chlorophyll concentrations and nitrogen across the vertical canopy profile affected by elevated CO_2_ in mature Quercus trees? Trees 36(6): 1797–1809. 10.1007/s00468-022-02328-7

15. Giolai M, Laine A-L. 2024. A trade-off between investment in molecular defense repertoires and growth in plants. Science 386(6722): 677–680. 10.1126/science.adn2779

16. Glazebrook J. 2005. Contrasting Mechanisms of Defense Against Biotrophic and Necrotrophic Pathogens. Annual Review of Phytopathology 43(Volume 43, 2005): 205–227. 10.1146/annurev.phyto.43.040204.135923

17. Hamilton JG, Zangerl AR, Berenbaum MR, Pippen J, Aldea M, DeLucia EH. 2004. Insect herbivory in an intact forest understory under experimental CO_2_ enrichment. Oecologia 138(4): 566–573. 10.1007/s00442-003-1463-5

18. Hart KM, Curioni G, Blaen P, Harper NJ, Miles P, Lewin KF, Nagy J, Bannister EJ, Cai XM, Thomas RM, et al. 2020. Characteristics of free air carbon dioxide enrichment of a northern temperate mature forest. Glob Chang Biol 26(2): 1023–1037. 10.1111/gcb.14786

19. IPCC 2023. Summary for policymakers. In: Lee HaR, J. ed. Climate Change 2023: Synthesis Report. Contribution of Working Groups I, II and III to the Sixth Assessment Report of the Intergovernmental Panel on Climate Change. Geneva, Switzerland: IPCC, 1–34.

20. Johnson SN, Waterman JM, Hall CR. 2020. Increased insect herbivore performance under elevated CO_2_ is associated with lower plant defence signalling and minimal declines in nutritional quality. Scientific Reports 10(1): 14553. 10.1038/s41598-020-70823-3

21. Kazan K. 2018. Plant-biotic interactions under elevated CO_2_: A molecular perspective. Environmental and Experimental Botany 153: 249–261. 10.1016/j.envexpbot.2018.06.005

22. Keeling RFK, Charles D. 2017. Atmospheric Monthly In Situ CO_2_ Data - Mauna Loa Observatory, Hawaii In Collections USDLD.

23. Kulman HM. 1971. Effects of Insect Defoliation on Growth and Mortality of Trees. Annual Review of Entomology 16: 289–324. 10.1146/annurev.en.16.010171.001445

24. Lake JA, Wade RN. 2009. Plant-pathogen interactions and elevated CO_2_: morphological changes in favour of pathogens. J Exp Bot 60(11): 3123–3131. 10.1093/jxb/erp147

25. Lan XT, P.; Thoning, K. W. 2025. Trends in globally-averaged CO_2_ determined from NOAA Global Monitoring Laboratory measurements: NOAA Global Monitoring Laboratory. 10.15138/9N0H-ZH07

26. Lessin R, Ghini R. 2009. Effect of increased atmospheric CO_2_ concentration on powdery mildew and growth of soybean plants. Tropical Plant Pathology 34: 385–392. 10.1590/S1982-56762009000600004

27. Lonsdale D. 2016. Powdery mildew of oak: a familiar sight with some hidden surprises. The ARB Magazine 43: 48–52.

28. MacKenzie AR, Krause S, Hart KM, Thomas RM, Blaen PJ, Hamilton RL, Curioni G, Quick SE, Kourmouli A, Hannah DM, et al. 2021. BIFoR FACE: Water–soil–vegetation–atmosphere data from a temperate deciduous forest catchment, including under elevated CO. Hydrological Processes 35(3): e14096. 10.1002/hyp.14096

29. Manresa-Grao M, Pastor-Fernández J, Sanchez-Bel P, Jaques JA, Pastor V, Flors V. 2022. Mycorrhizal Symbiosis Triggers Local Resistance in Citrus Plants Against Spider Mites. Frontiers in Plant Science Volume 13–2022. 10.3389/fpls.2022.867778

30. Marçais B, Desprez-Loustau M-L. 2014. European oak powdery mildew: impact on trees, effects of environmental factors, and potential effects of climate change. Annals of Forest Science 71(6): 633–642. 10.1007/s13595-012-0252-x

31. Marçais B, Kavkova M, Desprez-Loustau M-L. 2009. Phenotypic variation in the phenology of ascospore production between European populations of oak powdery mildew. Annals of Forest Science 66(8): 814–814. 10.1051/forest/2009077

32. Marçais B, Piou D, Dezette D, Desprez-Loustau M-L. 2017. Can Oak Powdery Mildew Severity be Explained by Indirect Effects of Climate on the Composition of the Erysiphe Pathogenic Complex? Phytopathology® 107(5): 570–579. 10.1094/phyto-07-16-0268-r

33. Maurino VG, Engqvist MK. 2015. 2-Hydroxy Acids in Plant Metabolism. Arabidopsis Book 13: e0182. 10.1199/tab.0182

34. Mayoral C, Ioni S, Luna E, Crowley LM, Hayward SAL, Sadler JP, MacKenzie AR. 2023. Elevated CO_2_ does not improve seedling performance in a naturally regenerated oak woodland exposed to biotic stressors. Frontiers in Forests and Global Change Volume 6–2023. 10.3389/ffgc.2023.1278409

35. McKown AD, Guy RD, Azam MS, Drewes EC, Quamme LK. 2013. Seasonality and phenology alter functional leaf traits. Oecologia 172(3): 653–665. 10.1007/s00442-012-2531-5

36. Nations FaAOotU. 2014. State of the World’s Forests 2014: Enhancing the Socioeconomic Benefits from Forests. Rome, Italy: Food and Agriculture Organization of the United Nations.

37. Norby RJ, Loader NJ, Mayoral C, Ullah S, Curioni G, Smith AR, Reay MK, van Wijngaarden K, Amjad MS, Brettle D, et al. 2024. Enhanced woody biomass production in a mature temperate forest under elevated CO_2_. Nature Climate Change 14(9): 983–988. 10.1038/s41558-024-02090-3

38. Norby RJ, Pastor J, Melillo JM. 1986. Carbon-nitrogen interactions in CO_2_-enriched white oak: physiological and long-term perspectives. Tree Physiology 2(1-2-3): 233–241. 10.1093/treephys/2.1-2-3.233

39. Office M 2023. Record breaking 2022 indicative of future UK climate: Met Office.

40. Özer N, Şabudak T, Özer C, Gindro K, Schnee S, Solak E. 2017. Investigations on the role of cuticular wax in resistance to powdery mildew in grapevine. Journal of General Plant Pathology 83(5): 316–328. 10.1007/s10327-017-0728-5

41. Pap P, Rankovic B, Masirevic S. 2013. Effect of temperature, relative humidity and light on conidial germination of oak powdery mildew (Microsphaera alphitoides Griff. et Maubl.) under controlled conditions. Archives of Biological Sciences 65: 1069–1077. 10.2298/ABS1303069P

42. Pétriacq P, Stassen JHM, Ton J. 2016. Spore Density Determines Infection Strategy by the Plant Pathogenic Fungus Plectosphaerella cucumerina Plant Physiology 170(4): 2325–2339. 10.1104/pp.15.00551

43. Pugliese M, Gullino ML, Garibaldi A. 2010. Effects of elevated CO_2_ and temperature on interactions of grapevine and powdery mildew: first results under phytotron conditions. Journal of Plant Diseases and Protection 117(1): 9–14. 10.1007/BF03356327

44. Quine C, Atkinson N, Denman S, Desprez-Loustau M, Jackson R, Kirby K. 2019. An assessment of the current evidence on oak health in the UK, identification of evidence gaps and prioritisation of research needs: Action Oak Knowledge review.

45. Rieske LK, Dillaway DN. 2008. Response of two oak species to extensive defoliation: Tree growth and vigor, phytochemistry, and herbivore suitability. Forest Ecology and Management 256(1): 121–128. 10.1016/j.foreco.2008.04.015

46. Roberts AJ, Crowley LM, Sadler JP, Nguyen TTT, Gardner AM, Hayward SAL, Metcalfe DB. 2022. Effects of Elevated Atmospheric CO_2_ Concentration on Insect Herbivory and Nutrient Fluxes in a Mature Temperate Forest. Forests 13(7): 998. 10.3390/f13070998

47. Robinson EAR, Geraldine D.; Newman, Jonathan A. 2012. A meta-analytical review of the effects of elevated CO_2_ on plant–arthropod interactions highlights the importance of interacting environmental and biological variables. New Phytol 194(2): 321–336. 10.1111/j.1469-8137.2012.04074.x

48. Roland JE, Douglas G. 1995. Biological control of the winter moth. Annual Review of Entomology 40: 475–492. 10.1146/annurev.en.40.010195.002355

49. Rumeau M, Sgouridis F, MacKenzie R, Carrillo Y, Reay MK, Hartley IP, Ullah S. 2024. The role of rhizosphere in enhancing N availability in a mature temperate forest under elevated CO_2_. Soil Biology and Biochemistry 197: 109537. 10.1016/j.soilbio.2024.109537

50. Ryan GD, Rasmussen S, Newman JA 2010. Global Atmospheric Change and Trophic Interactions: Are There Any General Responses? In: Baluška F, Ninkovic V eds. Plant Communication from an Ecological Perspective. Berlin, Heidelberg: Springer Berlin Heidelberg, 179–214.

51. Sanchez-Lucas R, Mayoral C, Raw M, Mousouraki M-A, Luna E. 2023. Elevated CO_2_ alters photosynthesis, growth and susceptibility to powdery mildew of oak seedlings. Biochemical Journal 480(17): 1429–1443. 10.1042/bcj20230002

52. Schausberger C, Roach T, Stöggl W, Arc E, Finch-Savage WE, Kranner I. 2019. Abscisic acid- determined seed vigour differences do not influence redox regulation during ageing. Biochem J 476(6): 965–974. 10.1042/bcj20180903

53. Schindelin J, Arganda-Carreras I, Frise E, Kaynig V, Longair M, Pietzsch T, Preibisch S, Rueden C, Saalfeld S, Schmid B, et al. 2012. Fiji: an open-source platform for biological-image analysis. Nature Methods 9(7): 676–682. 10.1038/nmeth.2019

54. Schymanski EL, Jeon J, Gulde R, Fenner K, Ruff M, Singer HP, Hollender J. 2014. Identifying Small Molecules via High Resolution Mass Spectrometry: Communicating Confidence. Environmental Science & Technology 48(4): 2097–2098. 10.1021/es5002105

55. Serrano M, Coluccia F, Torres M, L’Haridon F, Métraux J-P. 2014. The cuticle and plant defense to pathogens. Frontiers in Plant Science Volume 5–2014. 10.3389/fpls.2014.00274

56. Sgouridis F, Reay M, Cotchim S, Ma J, Radu A, Ullah S. 2023. Stimulation of soil gross nitrogen transformations and nitrous oxide emission under Free air CO_2_ enrichment in a mature temperate oak forest at BIFoR-FACE. Soil Biology and Biochemistry 184: 109072. 10.1016/j.soilbio.2023.109072

57. Skwarek-Fadecka M, Nawrocka J, Sieczyńska K, Patykowski J, Posmyk MM. 2024. Effect of Oak Powdery Mildew on Ascorbate–Glutathione Cycle and Other Antioxidants in Plant— Erysiphe alphitoides Interaction. Cells 13(12): 1035. 10.3390/cells13121035

58. Smith AG, Croft MT, Moulin M, Webb ME. 2007. Plants need their vitamins too. Current Opinion in Plant Biology 10(3): 266–275. 10.1016/j.pbi.2007.04.009

59. Smith CA, Want EJ, O’Maille G, Abagyan R, Siuzdak G. 2006. XCMS: Processing Mass Spectrometry Data for Metabolite Profiling Using Nonlinear Peak Alignment, Matching, and Identification. Analytical Chemistry 78(3): 779–787. 10.1021/ac051437y

60. Smith F, Luna E. 2023. Elevated atmospheric carbon dioxide and plant immunity to fungal pathogens: do the risks outweigh the benefits? Biochemical Journal 480(22): 1791–1804. 10.1042/bcj20230152

61. Song M, Peng L, Shang Y, Zhao X. 2022. Green technology progress and total factor productivity of resource-based enterprises: A perspective of technical compensation of environmental regulation. Technological Forecasting and Social Change 174: 121276. 10.1016/j.techfore.2021.121276

62. Teshome DT, Zharare GE, Naidoo S. 2020. The Threat of the Combined Effect of Biotic and Abiotic Stress Factors in Forestry Under a Changing Climate. Frontiers in Plant Science Volume 11–2020. 10.3389/fpls.2020.601009

63. Whittet R, Cavers S, Ennos R, Cottrell J. 2019. Genetic considerations for provenance choice of native trees under climate change in England. Genetic considerations for provenance choice of native trees under climate change in England: viii + 44 pp.

64. Williams A, Petriacq P, Schwarzenbacher RE, Beerling DJ, Ton J. 2018. Mechanisms of glacial- to-future atmospheric CO(2) effects on plant immunity. New Phytol 218(2): 752–761. 10.1111/nph.15018

