## Supplementary Figure 1 for "Multi-year study on the effects of elevated CO_2_ in mature oaks unravels subtle metabolic adjustments but stable biotic stress resistance"

| 0 | 1 | 2 | 3 | 4 | 5 |
|---|---|---|---|---|---|
| 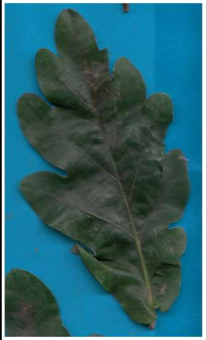 | 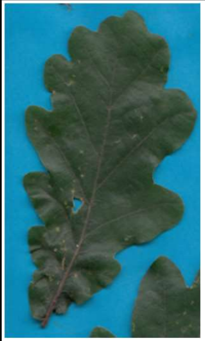 | 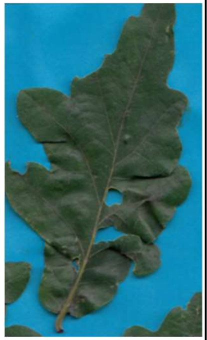 | 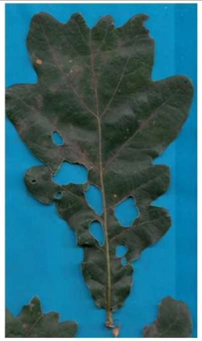 | 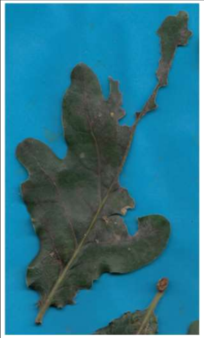 | 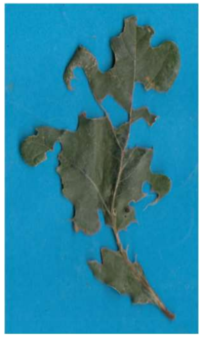 |
| Undamaged leaf | Small hole | Larger holes | Significant holes | Significant leaf damage | Near Complete/ Complete leaf damage |
| No damage 0% | Damage <10% | 10-25% damage | 26-50% damage | 51-75% damage | 75%< damage |

**Supplementary Figure 1. Representative images and scoring scale for assessment of insect damage.** First row represents the categories, second includes the representative images for each category, third represents the damage as observed in the images and forth represents the damage as a % of total leaf area affected.
