## Supplementary Figure 2 for "Multi-year study on the effects of elevated CO_2_ in mature oaks unravels subtle metabolic adjustments but stable biotic stress resistance"

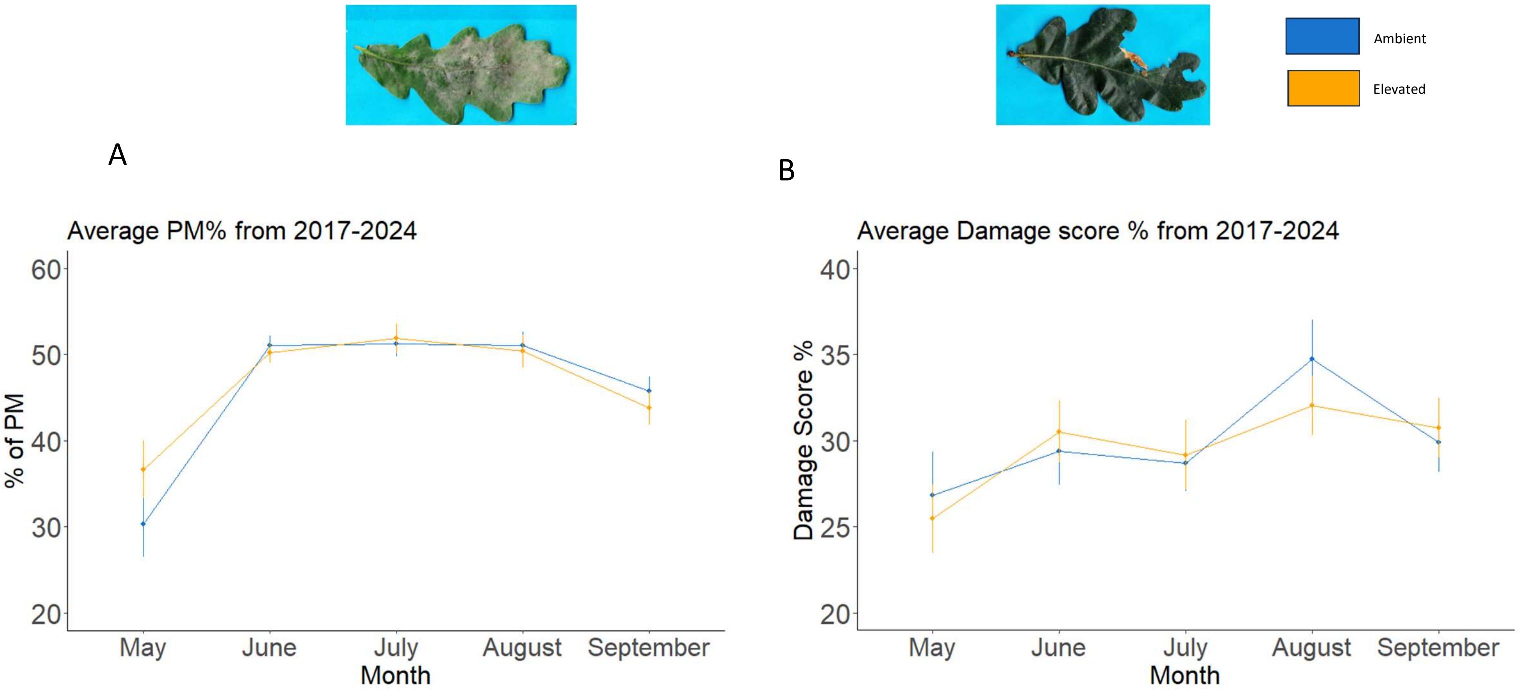


**Supplementary Figure 2. Growth season comparison between ambient $CO_2$ and elevated $CO_2$ treatments.** A) % of powdery mildew (PM) from May to September considering all years (i.e. from 2017 to 2024). B) % of insect damage score from May to September considering all years (i.e. from 2017 to 2024).
