## Supplementary Figure 3 for "Multi-year study on the effects of elevated CO_2_ in mature oaks unravels subtle metabolic adjustments but stable biotic stress resistance"

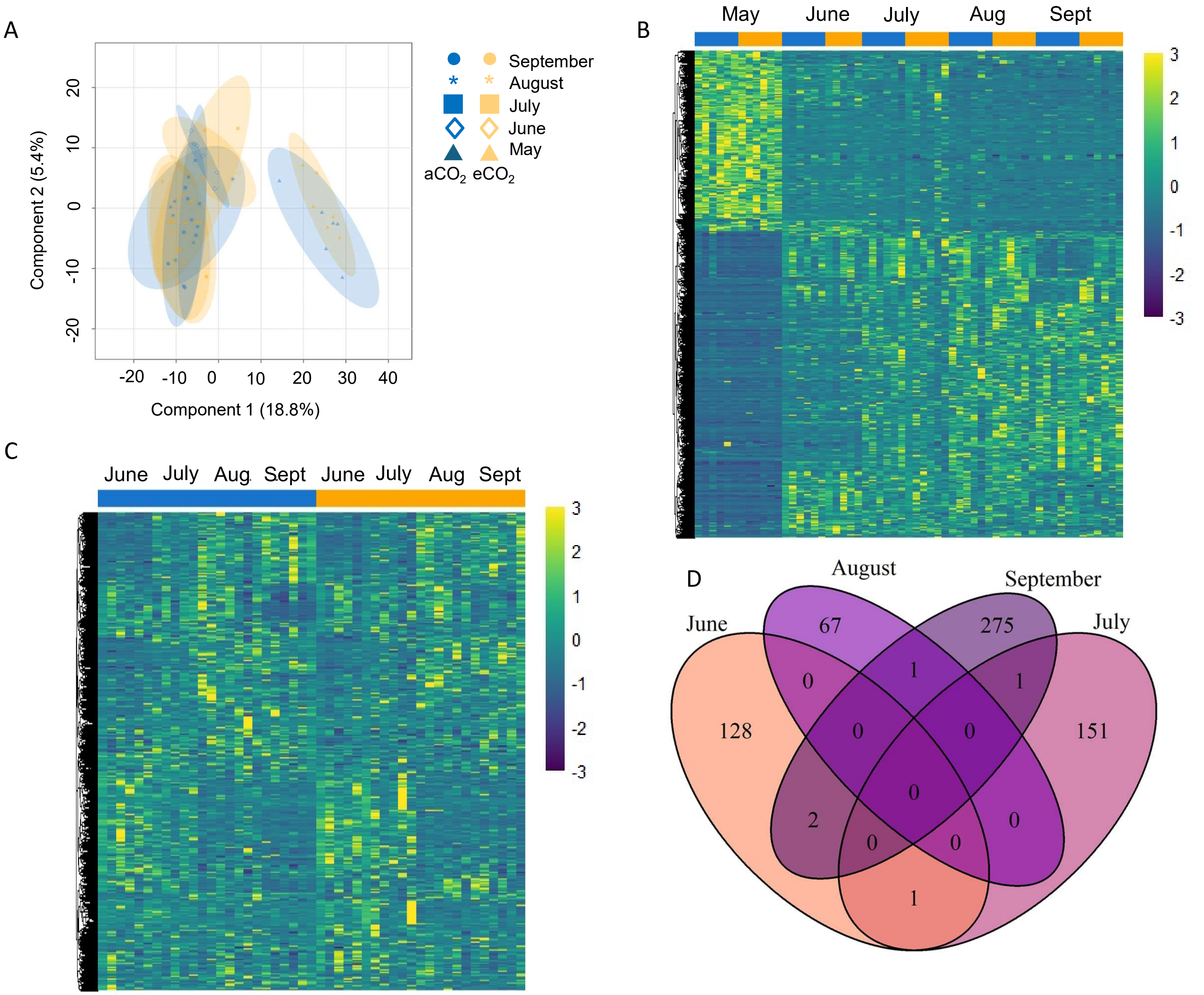


**Supplementary Figure 3. Global analysis of metabolomics data.** A) Combined principal component analysis (PCA) of metabolomic profiles across all sampling months and $CO_2$ treatments. PCA was performed on untargeted metabolomic data collected from oak leaves sampled monthly between May and September 2021 under ambient ($aCO_2$) and elevated $CO_2$ ($eCO_2$) conditions. B) Clustering analyses of differentially accumulated mass-to-charge (m/z) values across the 2021 growing season. Heatmap clustering of all differentially accumulated m/z features across May-September, grouped by both month and $CO_2$ treatment. C) Clustering analysis excluding May and combining ambient $CO_2$ ($aCO_2$) and elevated $CO_2$ ($eCO_2$) samples. D) Venn diagram of the overlap between differently accumulated m/z values under $aCO_2$ and $eCO_2$ in June, July, August and September.
